# Low-molecular-weight *Ulva lacinulata* extract exhibiting anti-inflammatory and pro-autophagic activities in RAW 264.7 macrophages: a promising candidate for the development of active ingredients targeting low-grade inflammation

**DOI:** 10.64898/2026.07.07.734444

**Authors:** J. Cherfan, D. Heerah, P-E. Bodet, Hania Haroune, B. Musnier, J. Saliba, R. Sulpice, J. Bodin, D. Dufour, X. Fioramonti, A.L. Dinel, C. Joffre, P. Delmarre, J. Le Faouder, E. Bouvret, I. Arnaudin, T. Maugard, N. Bridiau

**Affiliations:** UMR 7266 LIENSs (CNRS/La Rochelle Université); 2, rue Olympe de Gouges, La Rochelle, France; Plant Systems Biology lab; School of Chemical and Biological Sciences, Ryan Institute, University of Galway, Orbsen Building, H91DK59, Galway, Ireland; SEPROSYS (Séparations, Procédés, Systèmes); 12 Rue Marie-Aline Dusseau, 17000 La Rochelle, France; Université Bordeaux, INRAE, Bordeaux INP, NutriNeuro, UMR 1286; Bâtiment UFR de Pharmacie, 146 Rue Léo Saignat, 33076 Bordeaux, France; NutriBrain Research and Technology Transfer, NutriNeuro, 33076 Bordeaux, France; Nutridry; 8 Av. de la Gare, 33840 Captieux, France; Abyss Ingredients; 860 Rte de Caudan, 56850 Caudan, France

**Keywords:** *Ulva lacinulata* low-molecular weight extract, comprehensive composition, low-grade inflammation, anti-inflammatory activity, pro-autophagic activity, functional ingredient

## Abstract

Marine macroalgae are valuable sources of bioactive compounds. In this study, we thus investigated the chemical composition and biological activity of an extract from the green seaweed *Ulva lacinulata*, composed of small bioactive compounds. Comprehensive compositional analyses and high-resolution mass spectrometry revealed its diverse molecular profile composed in particular of peptides/amino acid derivatives, saccharides, low-chain fatty diacids, oxylipins and minerals. Its anti-inflammatory activity was assessed after 6 h pre-treatment in LPS-stimulated cultured RAW 264.7 macrophages, showing that it significantly and dose-dependently reduced the expression and/or secretion of pro-inflammatory cytokines such as TNF-α and IL-6, and targeted the NF-κB signaling cascade. It modulated the SIRT1-AMPK signaling axis and increased the LC3-II/LC3-I ratio, supporting the activation of a controlled autophagic response. This work highlighted the potential of this marine-derived extract as a safe and effective functional ingredient for the development of functional food and/or dietary supplements targeting chronic low-grade inflammation.

## Introduction

Marine macroalgae have been increasingly recognized as valuable sources of bioactive compounds for the development of functional food ingredients and dietary supplements^1^. In recent years, growing attention has indeed been paid to marine-derived resources as sustainable alternatives to terrestrial bioactive sources, driven by the need for novel ingredients combining health benefits with environmental resilience. In this context, green macroalgae such as species of the genus *Ulva* represent particularly promising candidates due to their broad ecological distribution, rapid growth rate, minimal cultivation requirements, and long history of safe consumption^2^. They also have attracted particular interest due to their capacity to synthesize a diverse array of metabolites—including amino acid derivatives, small carbohydrates, peptides, polyphenols, pigments, and sulfur-containing compounds—many of which have been associated with antioxidant, immunomodulatory, and anti-inflammatory activities^3^, highlighting their potential as multi-target bioactive compounds. However, *Ulva* biomass also contains substantial amounts of high–molecular weight (MW) polysaccharides, especially ulvans, which are complex highly sulfated polymers^4^. Although ulvans have been widely studied, several reports suggest that certain high–MW ulvan structures or sulfation patterns can stimulate innate immune pathways and enhance immunomodulatory and/or pro-inflammatory responses in macrophages^5–7^. This duality in biological activity depending on molecular size and structural features underscores the importance of refining extraction strategies to selectively enrich fractions with beneficial properties. For applications that require potent anti-inflammatory properties, selectively isolating low–MW metabolites from *Ulva* macroalgae—while excluding polysaccharides with potential immunostimulatory effects—is therefore essential. This is particularly relevant in the context of chronic low-grade inflammation, which is a persistent but subtle inflammatory state now widely recognized as a central driver of metabolic dysregulation, and a major contributing factor to the development of insulin resistance, cardiovascular diseases, intestinal imbalance, and functional decline associated to various physio pathological contexts, such as aging or overtraining^8–10^.

Unlike acute inflammatory responses, this persistent and subclinical inflammatory state is characterized by sustained activation of innate immune pathways and low-level production of pro-inflammatory mediators, which progressively alter metabolic homeostasis and tissue function^11^. At the cellular level, macrophages play a central role in this process by integrating environmental and metabolic cues and orchestrating the production of inflammatory mediators, making them a key target for interventions aimed at restoring immune balance^12^. Consequently, natural compounds capable of attenuating pro-inflammatory signaling in macrophages and modulating systemic immune tone hold strong potential for the development of dietary supplements aimed at long-term inflammation control^13^. By targeting upstream inflammatory regulators and supporting cellular adaptive responses rather than inducing immunosuppression, such nutritional strategies may contribute to the maintenance of metabolic resilience and overall health during ageing and in populations exposed to chronic inflammatory stress.

In this context, the present study investigated a low–MW extract from the marine macroalga *Ulva lacinulata* produced through a patented industrial process enabling the selective recovery of small bioactive molecules while excluding large biopolymers such as polysaccharides. This extract was chemically characterized using proximate analyses and high-resolution mass spectrometry to obtain its detailed molecular profile and then evaluated regarding its anti-inflammatory capacity in RAW 264.7 macrophage models under LPS-induced inflammatory stimulation. Beyond assessing its capacity to modulate inflammatory responses, a key aim of the study was to explore its potential mechanisms of action, with a particular focus on autophagy induction, a cellular process increasingly recognized for its ability to regulate inflammatory signaling. The interplay between immuno-metabolic regulators and autophagic pathways has indeed emerged as a critical determinant of macrophage functional plasticity and inflammatory outcomes^14^. This integrated compositional and cellular investigation aimed to determine whether this extract could exhibit anti-inflammatory properties consistent with its further development as an innovative active ingredient for functional food and dietary supplements targeting chronic low-grade inflammatory conditions.

## Materials and Methods

### Materials

Phosphate-buffered saline (PBS), radio immunoprecipitation assay (RIPA) lysis buffer, protease/phosphatase inhibitor cocktail, bicinchoninic acid (BCA) protein assay kit and enhanced chemiluminescence (ECL) substrate were purchased from Bio-Rad (Marnes-La-Coquette, France). Plates and microplates for cell culture were purchased from Bioswisstec (BD Falcon, Schaffhausen, Switzerland). Lipopolysaccharide (LPS) from *Escherichia coli* 0111:B4 and all chemicals used were from Merck (Sigma-Aldrich Chimie, Saint-Quentin Fallavier, France), unless otherwise stated in the text.

### Methods

#### Preparation of the *Ulva lacinulata* extract, so-called “permeate”

*Ulva lacinulata* permeate was produced using a patented industrial extraction and fractionation designed to selectively recover low-molecular-weight compounds^15,16^, slightly modified.

Fresh collected algae were first washed with sea water to remove impurities and steeped at room temperature (RT), in a tap water bath for less than 15 min, to strongly limit molecular diffusion. The partially desalted algae were then wrung using a wringing system and air-dried at less than 40°C. Once dried, they were finally ground using a grinder until obtaining algal particle sizes within the range 500-1000 µm. A total of 500 g of *Ulva lacinulata* dried flakes was then transferred into a thermostated tank at 80°C containing 2.5 l of distilled water. The aqueous extraction was made under constant agitation with a bladed stirrer at a rotation speed of 10 spins/min for 2 h. The pulps were then removed from the tank and filtered with the fabric cone to collect the aqueous extract. Then, 13 l of extract was recovered and filtered with an ultrafiltration (UF) unit equipped with a 15kDa Kerasep KBW membrane (Novasep Process, Pompey, France). The filtration was carried out at 80°C at a pressure of 5 bar and a circulation flow of 450 L/h (circulation speed of 5 m/s) until obtaining a retentate around 4°B. Subsequently, a major part of algal-derived proteins and polysaccharides were transferred into the UF retentate and thus separated from the UF permeate^16,17^. Finally, 1.5L of the UF permeate was concentrated by reverse osmosis at less than 30°C, until measuring more than 10°B, frozen at −20°C and then −80°C, freeze-dried using a Heto PowerDry OL 6000 lyophilization system (Thermo Electron Co., France) and ground into a fine powder using a cooking blender for approximately 30 s, to obtain a white cream-colored solid powder, designated as “permeate” within the text and always kept at 20°C under argon atmosphere to prevent it from rehydration and/or oxidation.

For all cell assays, a stock solution of permeate (1 mg/ml) was prepared in Dulbecco’s Modified Eagle Medium (DMEM, PAN Biotech, Aidenbach, Germany), sterilised through 0.22 μm filtration and immediately used.

#### Preparation of *Ulva lacinulata* permeate fractions

Three different sample pre-treatment procedures were applied to obtain more simple and enriched fractions starting from permeate: 1) a size-based fractionation method that was also used to determine the weight distribution of permeate (Method 1; Figure S1); 2) three different forms of differential precipitation procedures associated to filtration methods (methods 2A, 2B and 2C; Figures S2, S3 and S4); and 3) two forms of acidic treatment using supported proton exchange. Acidic treatment via supported proton exchange was conducted by using a strong acidic cation-exchange resin (Amberlyst™ 15 dry; 4.7 meq H^+^/g_dry material_). Briefly, 20 ml of permeate solution at 25 mg/ml was placed in a 50 ml bottle already containing 5 g of Amberlyst™ 15 dry resin previously hydrated and washed with ultrapure water. This batch system was then hermetically closed and placed in an LSE cabinet-style shaking incubator (Corning, NY, USA), maintained at 60°C under orbital shaking at 120 rpm. Then, 1.5 ml aliquots were collected after 30 min and 5 h, frozen in an ice bath and neutralized at pH 7.0 by adding 1 M NaOH. They were finally submitted to 0.22 µm filtration prior to freeze-drying, leading to the production of permeate fractions 3a (30 min treatment) and 3b (5 h treatment).

All permeate solid fractions were obtained by freeze-drying at −20°C and then −80°C, using a Heto PowerDry OL 6000 lyophilization system (Thermo Electron Co., France).

#### Biochemical composition

The total sugar content was determined according to the phenol-sulfuric method^18^, using rhamnose as a standard. Neutral sugars and uronic acids (acidic sugars) content determination was carried out using the resorcinol method of Monsigny et al.^19^ and m-hydroxydiphenyl (mHDP) method of Blumenkrantz et al.^20^, respectively. Sample concentrations in neutral sugars (NS) and uronic acids (UA) were calculated according to the following equations, adapted from Montreuil et Spik^21^ and Filisetti-Cozzi and Carpita (1991)^22^, to avoid reciprocal interferences between these two methods:

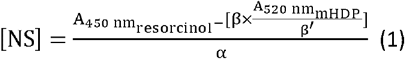

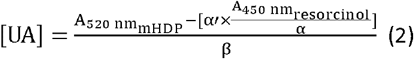

with:

α: calibration slope obtained with the resorcinol analysis of glucose, used as a neutral sugar standard

α′: calibration slope obtained with the mHDP analysis of glucose, used as a neutral sugar standard

β: calibration slope obtained with the resorcinol analysis of glucuronic acid, used as an uronic acid standard

β′: calibration slope obtained with the mHDP analysis of glucuronic acid, used as an uronic acid standard

The osidic composition was determined by the CEVA laboratory (Pleubian, France)^23^, using reverse-phase HPLC analysis after depolymerization under methanol-acid hydrolysis reaction (methanolysis)^24,25^. Starch content was subsequently extrapolated from total glucose content according to the following equation:

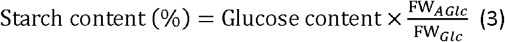

With:

Glucose content: 2.4%

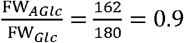: molecular weight ratio of polymeric anhydroglucose over monomeric glucose

Two methods were used to assess the sulfate group content of oligo- and poly-saccharides. Inorganic and organic sulfate contents were determined using the turbidimetric method, as described by Dodgson and Price^26^, adapted and optimized for algae polysaccharide-rich samples by Torres et al. (2021)^27^. Given that the sample studied in this work was supposed to be mainly composed of oligosaccharides and/or low MW polysaccharides, its polysaccharide specific sulfate content was estimated using 3-amino-7-(dimethylamino)phenothizin-5-ium chloride (Azure A), which binds the sulfate groups in a polysaccharide chain^28,29^, using dextran sulfate (17% sulfur) as a standard.

The quantification method of polyphenols was adapted from the original one^30^, using gallic acid as a standard^31^.

The protein content was determined using the Kjeldahl method (total nitrogen × 5.0)^32^. The amino acid composition, tryptophan excepted, and the total tryptophan content were determined by the Upscience laboratory (56250 SAINT-NOLFF, France)^33^, according to internal methodologies (ACIDAM 96 and TRYPRO 95) involving total acidic or alkaline hydrolysis, respectively, followed by ion HPLC-UV analysis with post-column derivation.

The lipid content was measured via the method of Chabrol and Charonnat^34^, using vegetal oil as a standard.

The ash content was determined by measuring the mass loss of samples heated for 4 h at 550°C.

#### Mineral composition

Total Hg analysis was carried out with an Advanced Mercury Analyser Altec AMA 254 (Symalab, Morlaàs, France), on dried sample powder aliquots ranging from 50 to 220 mg, weighed to the nearest 0.01 mg. The other elements were analysed using an Agilent 5800VDV ICP-AES and an Agilent 8900 QQQ ICP-MS (Agilent Technologies, Les Ulis, France), after acidic digestion: around 250 mg of sample was weighed and then digested using a 67–70% HNO_3_/34–37% HCl (6:2, v/v) mixture (trace metal quality, Thermo Fisher Scientific, Illkirch-Graffenstaden, France), carried out overnight at room temperature and then in an ETHOS Up microwave oven (Milestone, Sorisole, Italy), firstly for 30 min using constantly increasing temperature up to 120°C, and secondly for15 min at 120°C. Each sample was made up to 50 mL with ultrapure quality water. Certified reference materials (lobster hepatopancreas TORT-3 and dogfish liver DOLT-5; National Research Council, Canada) were prepared and analysed under the same conditions as the samples.

#### Compounds identification by ultra-high pressure liquid chromatography coupled and high-resolution mass spectrometry (UHPLC-HRMS)

UHPLC-HRMS analyses were performed with an Acquity UHPLC H-class system (Waters, Milford, USA) coupled to a XEVO G2S Q-Tof mass spectrometer equipped with an electrospray ionization source (Waters, Manchester, England). For injection, all dried permeate fractions were solubilized at 1 mg/ml in water/formic acid (100:0.1, v/v) and/or in an acetonitrile/water/formic acid (30:70:0.1, v/v/v), centrifuged at 13.000 rpm for 5 min, and filtered through a 0.22 μm PVDF filter. Then, 10 μL of the obtained sample was injected into a BEH C_18_ column (2.1 x 150 mm; 1.7 μm) at an analysis temperature maintained at 40°C. Separation of compounds was obtained using a gradient method involving volume mixtures of solvents water/formic acid (100:0.1, v/v) (**solvent A**) and acetonitrile/formic acid (100:0.1, v/v/v) (**solvent B**) as follows: 0-3 min, 70% A and 30% B; 3-4 min 20% A and 80% B; 4-6 min 20% A and 80% B; 6-6.5 min, 0% A and 100% B; 6.5-10 min 0% A and 100% B; 10-10.1 min, 70% A and 30% B; 10.1-15 min, 70% A and 30% B. Mass spectrometry data were acquired in both polarities (capillary voltages of +2,5 kV and −2 kV, respectively) using the following ESI-QToF setup: m/z within the range 50-2000; scan time of 0.1 s; sampling cone voltage of 35 V; source and desolvation gas temperatures of 120 and 500°C, respectively; cone gas and desolvation gas flows of 50 and 800 L/h, respectively. MS^2^ analyses were based on an ion intensity threshold of 500,000.s^−1^ (Data Dependent Analysis) with a collision energy ramping from 10 to 50 eV. All experiments were conducted with leucine enkephalin at 1 ng.µl^−1^ used as a lock mass to prevent mass deviations.

These parameters provided high-resolution separation and detection, enabling detailed identification and characterization of compounds in the permeate fractions. To do so, the resulting MS^1^ analyses were firstly used to conduct peak picking on the Workflow4metabolomics platform (W4m)^35,36^, with a signal/noise threshold set at 10. Then, retention times and m/z values were aligned and a list of detected ions for each polarity was thus obtained. In parallel, MS/MS (MS^2^) ion spectra were processed using the identification software Sirius4^©21^. Molecular formula determination was performed in Sirius, allowing the elements C, H, O, N, P, S, F, Cl, Br and I with an MS^2^ mass accuracy set to 10 ppm. These elements were considered with H^+^, Na^+^ and K^+^ as possible adducts in positive polarity and both Br^−^ or Cl^−^ and deprotonation ([M-H]^−^) as possible adducts in negative polarity. The ‘Predict FPs’ module was then applied to infer molecular fingerprints, while ‘Search DBs’ compared these predicted fingerprints to reference MS^2^ data from the 21 proposed databases including natural compounds databases such as ‘Human Metabolome Database (HMDB), ‘Chemical Entities of Biological Interest’ (ChEBI), ‘COlleCtion of Open NatUral producTs’ (COCONUT), and ‘Global Natural Products Social Molecular Networking’ (GNPS) repository, among others. Finally, the ‘CANOPUS’ module was used to predict compound classes. Molecular formulas were retained when “CSI:FingerID” scores representing the similarity between the candidate and the fragmentation profile exceeded −75, and only compound structures showing a minimum similarity of 60 % were considered for annotation. At the end, the list of ions obtained from W4M was completed by identification data from Sirius^©^.

Sixteen identified compounds were then directly detected in permeate, confirming their presence: abromine, arginyl-glutamine, azelaic acid, D-glucurono-6,3-lactone, gynesine, itaconate, stachydrine, suberate, sulfosalicylate and vitamin B5, 9-hydroxy-11-(3-hydroxy-5-(1-hydroxypropyl)tetrahydrofuran-2-yl)undec-10-enoic acid), alanyl-glutamyl-arginin, arginyl-pyroglutamate, citric acid, fulgidic acid and phytoprostane A1. To do so, a 1 mg/ml permeate solution was submitted to the same UHPLC-HRMS analysis protocol. Finally, pure standard solutions of abromine, arginyl-glutamine, azelaic acid, D-glucurono-6,3-lactone, gynesine, itaconate, stachydrine, suberate, sulfosalicylate and vitamin B5 within the calibration range 0.001-5 µg/ml were used to both strictly confirm the presence of these compounds in permeate by combining their retention time and MS^1^/MS^2^ data, and quantify them in 3 independent 1 mg/ml permeate solution, using the same UHPLC-HRMS analysis conditions.

#### Cell culture

RAW 264.7 murine macrophages were cultured in DMEM supplemented with 10% (v/v) fetal bovine serum and 1% (v/v) antibiotic solution (10,000 U/ml penicillin, 10 mg/ml streptomycin) (both from Gibco, Thermo Fisher Scientific, Illkirch-Graffenstaden, France), used as the complete culture medium, in a temperature-controlled humidified incubator with 5% CO2 at 37⍰C. Unless otherwise stated, cells at 1.10^6^ cells/ml in 1 ml of complete culture medium were seeded in 6-well plates and allowed to adhere overnight. The medium was then removed and 1 ml of permeate (0.1–100 µg/ml) was added for 6 h, followed by the incubation with LPS (1 µg/ml) for the next 18 h. Controls included unstimulated cells (basal conditions), cells incubated with LPS only, and cells preincubated with a well-known anti-inflammatory drug, dexamethasone, used at 1 µM as a positive anti-inflammatory control. A concentration of LPS of 1 µg/ml was selected for this study, as it is considered relatively standard in the literature^38^. Dexamethasone exhibits potent anti-inflammatory effects by suppressing LPS-induced responses and is widely used to treat inflammatory diseases^39^. By including these controls in our experimental design, we ensured that the macrophage responses were consistent with those reported in established literature models.

Mycoplasm level was checked in every case using the MycoAlert Mycoplasma Detection kit from Lonza (Basel, Switzerland) to ensure that cells were mycoplasma-free.

#### Cell viability assay

3-(4,5-dimethylthiazol-2-yl)-2,5-diphenyltetrazolium bromide (MTT) assay was used to assess the effect of permeate on cell viability. Briefly, the RAW 264.7 cells at 5.10^4^ cells/ml in 100 µl of complete culture medium were seeded in 96-well microplates and allowed to adhere for 24 h. The medium was then removed and 100 µl of permeate (0.1–250 µg/ml) or LPS (1 µg/ml) prepared in complete culture medium or negative control (complete culture medium) was added into the wells. After 24 h, 10 µl of MTT (5% (w/v) in PBS) was added into each well and the plates were incubated for 4 h at 37⍰C. The medium was then removed and 150 µl of dimethyl sulfoxide was added into each well to dissolve the MTT crystals, prior to incubation for 10 min and absorbance reading at 570 nm using a Fluostar Omega microplate reader (BMG Labtech, Champigny-sur-Marne, France).

### RT-qPCR

The anti-inflammatory effect of permeate on gene expression by RT-qPCR was carried out using a kinetic approach: after 24 h pre-exposure to 100 µg/ml permeate or 1 µM dexamethasone, cells were stimulated with the addition of LPS (1 µg/ml) and harvested for analysis after the next 2 h, 6 h, or 24 h.

Total RNA from RAW 264.7 macrophages was extracted using the TRIzol extraction protocol (Invitrogen, Thermo Fisher Scientific, Illkirch-Graffenstaden, France). The quantity of the extracted RNA was assessed using a Nanodrop spectrophotometer (Nanodrop One, Life Technologies, France). Two micrograms of RNA were submitted to reverse transcription to generate complementary DNA (cDNA) using Superscript IV (Invitrogen, Thermo Fisher Scientific, France). Subsequently, the cDNAs were amplified through PCR with TaqMan® primers specific to the target genes of TNFα (Mm00443258_m1), IL-6 (Mm00446190_m1) and IL-1β (Mm00434228_m1), as described by Mougin et al.^40^. B2m (Mm900437762_m1) was used as a reference gene. Fluorescence levels were assessed using a LightCycler® 480 instrument II (Roche, La Rochelle, France), and the final quantification was performed using the comparative threshold cycle (Ct) method. The outcomes were presented as relative expression levels, calculated in reference to the control target mRNA expression, as outlined by Chataigner et al.^41^.

#### Nitric Oxide Secretion (Griess Assay)

The RAW 264.7 cells at 4.10^5^ cells/ml in 100 µl of complete culture medium were seeded in 96-well microplates and allowed to adhere for 24 h. After 6 h permeate pre-treatment followed by 18 h LPS stimulation, culture supernatants were collected and centrifuged (1,000×g, 5 min). Equal volumes (100µl) of supernatant and Griess reagent (Invitrogen, Thermo Fisher Scientific, Illkirch-Graffenstaden, France) were mixed in a 96-well plate and incubated for 10 min at RT. Absorbance was read at 540 nm using a Fluostar Omega microplate reader (BMG Labtech, Champigny-sur-Marne, France). Secreted nitric oxide concentration (µM) was interpolated from a sodium nitric oxide calibration curve.

#### Cytokine Quantification (ELISA)

TNF-α, and IL-6 levels were measured in cell clarified supernatants using species-specific sandwich ELISA kits according to manufacturer’s instructions (Invitrogen, Thermo Fisher Scientific, Illkirch-Graffenstaden, France). Absorbance was read at 450 nm using a Fluostar Omega microplate reader (BMG Labtech, Champigny-sur-Marne, France). Concentrations in TNF-α and IL-6 were interpolated from calibration curves (four-parameter logistic fit).

#### Western blot analyses

Cells were seeded at a density of 1×10^6^ cells in 1 ml of complete culture medium in a 6 well microplate and allowed to adhere for 24 h. Proteins from cells were extracted using the RIPA lysis buffer supplemented with 1X protease/phosphatase inhibitor cocktail and quantified using the BCA protein assay kit from Bio-Rad. Equal amounts (20 µg) of proteins were mixed with 4X loading buffer (Bio-Rad, Marnes-La-Coquette, France) and then loaded on 4-20% pre-casted gels and transferred onto PVDF membranes using the transblot system from Bio-Rad (Marnes-La-Coquette, France). Membranes were blocked with 5% (w/v) half-fat milk solution for 1 h in Tris Buffered Saline containing 0.1% (v/v) Tween 20 (TBS-T, Bio-Rad, Marnes-La-Coquette, France) at RT, and then incubated overnight at 4°C with the following antibodies at 1/1000 dilution towards Cox2 (D5H5) XP Rabbit mAb, NLRP3 (D4D8T) Rabbit mAb, p-NFkB p65 (Ser536)(93H1) Rabbit mAb, NF-kB (D14E12) XP Rabbit mAb, SIRT1 (D1D7) Rabbit mAb, Bcl-2 (D17C4) Rabbit mAb, Phospho-AMPKα (Thr172) (40H9) Rabbit mAb, AMPKα (D5A2) Rabbit mAb, β-Actin (13E5) Rabbit mAb (purchased from Cell Signaling Technology, Ozyme, Saint-Cyr L’Ecole, France, except LC3 Polyclonal antibody that was from Proteintech, Thermo Fisher Scientific, Illkirch-Graffenstaden, France). Membranes were washed three times with TBS-T and then incubated with the appropriate horseradish peroxidase-conjugated secondary IgG antibody (1/1000 to 1/5000) at RT for 1.5 h. Immunodetection was performed using the Clarity western enhanced chemiluminescence kit (Bio-Rad, Marnes-La-Coquette, France) with a Chemidoc Imaging System (Bio-Rad, Marnes-La-Coquette, France). Optical densities were measured using ImageLab software (Bio-Rad, Marnes-La-Coquette, France). All optical density plots represent values normalized to loading controls, as indicated in the figure captions.

#### Confocal microscopy

The cells underwent 2–3 washes with PBS before being fixed with 1 ml of 4% (w/v) paraformaldehyde for 1 h at RT. After 3 additional washes with PBS, 1 ml of 1% (v/v) Triton X-100 in PBS was added to permeabilize the cells for 1 h at RT.

Regarding NF-κB nuclear translocation following, 3 washes with PBS were then applied prior to the addition of 100 μL of immunostaining blocking solution (5% (w/v) bovine serum albumin (BSA) in PBS), and incubation at 37°C for 1 h. After removal of the blocking solution, the cells were incubated overnight with 50 µl NF-κB p65 antibody solution (1/300 dilution, Cell Signaling Technology, Danvers, MA, USA), in a dark, humid chamber at 4°C. After another round of 2-3 PBS washes, 50 µl of immunofluorescence secondary antibody solution (Alexa Fluor 488-coupled secondary antibodies used at a dilution of 1/300, Cell Signaling Technology, Ozyme, Saint-Cyr L’Ecole, France) was added, and the cells were incubated in a dark, humid chamber at 37°C for 1 h. The cells were then treated with one droplet of fluorescence quenching solution containing 4⍰,6-diamidino-2-phenylindole (DAPI, Thermo Fisher Scientific, Illkirch-Graffenstaden, France). Coverslips were mounted onto glass slides using an antifade mounting medium.

Regarding cytoskeleton staining and fluorescence imaging, the cells were then incubated overnight in the dark with fluorescent Alexa Fluor 594 dye-conjugated phalloidin (Thermo Fisher Scientific, Illkirch-Graffenstaden, France), diluted in PBS containing 1% (m/v) BSA. After staining, the cells were washed with PBS and nuclei were counterstained with DAPI (Thermo Fisher Scientific, Illkirch-Graffenstaden, France) for 5 min. Coverslips were mounted onto glass slides using an antifade mounting medium.

Image acquisition was performed using an Axio Observer 7 microscope with an LSM 900 confocal head (lasers 405 nm, 488 nm and 561 nm, ZEISS, Oberkochen, Germany). In order to quantify the proportion of NF-κB translocated into the cell nucleus, the images were analyzed using custom-made macros with the open source Fiji/ImageJ software^42^ and the cytoplasmic NF-κB signal was subtracted from its nuclear signal, as described by Trask et al.^43^. The cell relative volume of macrophages was determined using the same software by measuring the total fluorescent Alexa Fluor 594 dye-conjugated phalloidin-positive cytoplasmic area and normalizing it to the number of nuclei detected in the corresponding field.

#### Statistical Analyses

Data analysis was performed using GraphPad Prism 8.0 (GraphPad software, 20 La Jolla, CA, USA). All data were presented as mean values ± SEM, of n independent experiments. The statistical significance of experiments involving multiple experimental groups compared to a single reference condition (Control or LPS-treated cells), was evaluated using a one-way analysis of variance (ANOVA), followed by a Dunnett’s multiple comparisons post-hoc test. This test was specifically selected to assess differences between each treatment condition and its corresponding control while controlling for multiple comparisons. A p-value <0.05 was considered statistically significant. Data were analyzed using parametric tests, assuming approximate normal distribution of residuals.

## Results

### Biochemical composition of Permeate

To have a broad view of the composition of permeate, both its weight distribution (Figure 1) and biochemical composition (Table 1) were estimated, using a method of size-based fractionation involving successive centrifugal ultrafiltration steps with different decreasing cut-off values on the one hand, and classical (bio-)chemical methods for the other.

**Table 1:** Biochemical composition of permeate.

| Content (% dw) |  |  |  |  |  |  |  |  |  |  |
| --- | --- | --- | --- | --- | --- | --- | --- | --- | --- | --- |
| Total sugars | Neutral sugars | Acidic sugars | Saccharide sulfates | Phenolics | Proteins / total nitrogen | Total amino acids | Lipids | Organic sulfates | Inorganic sulfates | Ash |
| 5.6 ± 0.3<br>(starch 2.2% <sup>a</sup>) | 5.0<br>± 0.7 | 5.6<br>± 0.5 | 0.14<br>± 0.007 | 0.45<br>± 0.001 | 24.6 ± 0.6 /<br>4.92 ± 0.12 | 17.6<br>± 1.4 | 0.54<br>± 0.017 | 1.01<br>± 0.37 | 2.45<br>± 0.20 | 51.0<br>± 1.0 |

<sup>a</sup> Obtained by extrapolation based on the total glucose content determination.

**Table 2:** Mineral composition of permeate.

| Content (mg/100 g dw) |  |  |  |  |  |  |  |  |  |  |  |  |
| --- | --- | --- | --- | --- | --- | --- | --- | --- | --- | --- | --- | --- |
| Na | Mg | K | Ca | P | S | I | Fe | Mn | Se | Zn | Cu | Mo |
| 1198.3<br>± 3.7 | 3265.5<br>± 42.0 | 7305.9<br>± 34.1 | 1764.7<br>± 17.1 | 595.0<br>± 1.5 | 10204.4<br>± 114.0 | 3.20<br>± 0.64 | 0.99<br>± 0.017 | 17.96<br>± 0.05 | 0.039<br>± 0.026 | 0.46<br>± 0.13 | 0.84<br>± 0.003 | 0.081<br>± 0.0019 |

**Figure 1:**
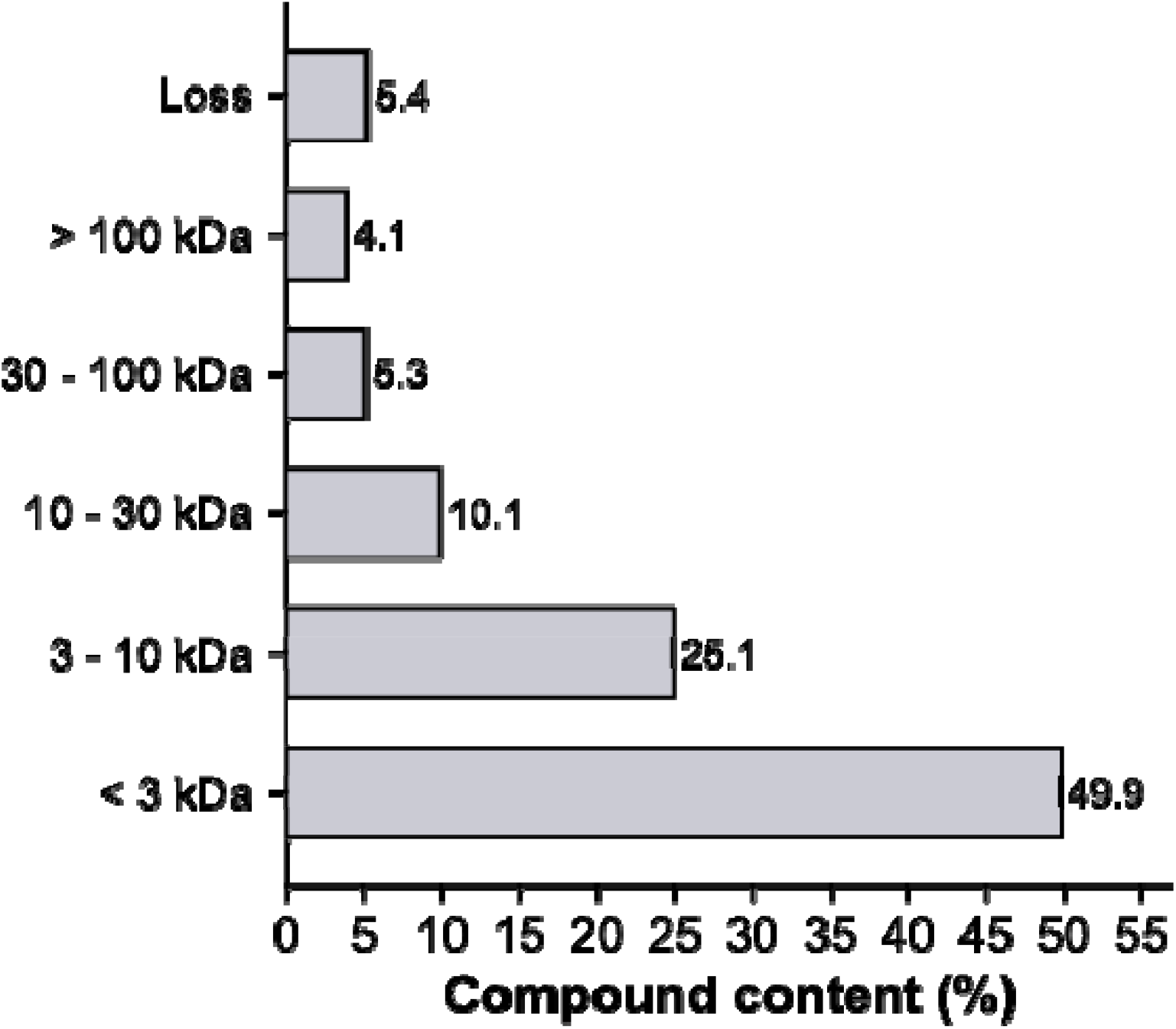
Weight distribution of compounds in permeate.

The MW distribution of permeate obtained by size-based fractionation logically indicated that it was mainly composed of low MW compounds (Figure 1): about 50% at the small compound and/or oligomer scale (lower than 3 kDa) and 25% at the oligomer or low MW polymer scale (MW values within the range 3-10 kDa). The 25% remaining was found to be constituted of 10% small polymers, comprised between 10 and 30 kDa, which can be explained by the 15 kDa cut-off value used to obtain permeate. Indeed, polymers of MW values within the range 10-15 kDa are expected to be found in this fraction. Moreover, this also statistically indicates that 10% at most of a whole population of compounds of less than 15 kDa are supposed to be retained by the UF membrane of such cut-off value^44^, which could lead to a slight overestimation of this 10-30 kDa fraction and should be considered as it would give a very similar result. In addition, it would explain why about 10% of the whole sample was found in the last fractions corresponding to cut-off values of 30 and 100 kDa, respectively.

The (bio-)chemical analyses of permeate confirmed and extended these preliminary results (Table 1), firstly confirming that permeate contained small compounds including a low content of lipids (0.54 ± 0.017% dw) and a high content of minerals (51% dw). Minerals were especially composed of inorganic sulfate (about 2.5% dw) and some macro (sodium, magnesium, potassium, calcium, phosphorus, sulfur) and trace elements (iodine, iron, manganese, selenium, zinc, copper, molybdenum) (Table 1). Seventeen heavy metals were analyzed at the same time (Pb, Ag, Cd, Hg, inorganic As, Ba, Co, Cr, Ni, Sn, V, Li, In, Al, Si, Sr, Ti; data not shown). All of them were far below the upper safe levels of minerals^45^, or heavy metals^46^, proving the safety of permeate regarding these elements.

It was also constituted of various bio(oligo)-polymers, proteins in particular (about 25%) and, to a lesser extent, saccharides: about 5-10% dw depending on the quantification method (about 5-6% dw when quantifying total sugars and 10-11% dw when adding neutral and acidic sugars). These differences could be mostly due to response variations depending on the monosaccharides used as standards, which were different in these quantification assays: Glc for total sugar determination; Glc for neutral sugars; GlcA for uronic acids. The osidic composition of permeate indicated that these saccharides were composed of various sugar moieties (Figure 2A), mainly neutral sugars, glucose and galactose in particular (45% Glc, 24% Gal, 9% Man, 8% Rha for the most represented) and a small content of uronic acids (<2% GlcA, GalA or IdoA). This composition is consistent with some of the most distributed polysaccharides in green seaweeds that are mainly glucans as storage polysaccharides and cellulose, arabinogalactans and rhamnans as parietal polysaccharides. Regarding seaweeds from the *Ulva* genus, rhamnans have been shown to be very typical and so-called ulvans, composed of ulvanobioruronic acid repeated disaccharide units A and B constituted of either glucuronic acid (about 90%) or iduronic acid (about 10%), respectively, and rhamnose, and associated to various minor sugar moieties such as glucose or xylose. Given that ulvans are sulphated polysaccharides, this was also in accordance with the determined organic (about 1% dw) and saccharide (about 0.15%) content values that indicated the presence in permeate of sulphated organic compounds including saccharides. Altogether, these results confirmed that permeate contained some glucan-type and rhamnan-type oligo- or poly-saccharides (likely starch and ulvan, respectively). Starch content was estimated approximately at 2.2% based on glucose content of permeate. The nature of (poly-)peptides and free amino acids was also explored by analysing the total amino acid composition of permeate (Figure 2B), which revealed that glycine, alanine, proline, lysine, and especially arginine (3.4% dw), aspartic acid (4.0% dw) and glutamic acid (5.5% dw) were the most represented amino acids in the totum. Besides, essential amino acids Val, Lys, Phe and Thr were significantly represented, within the range 0.15-0.51% dw. Permeate was also composed of a small but significant part of phenolic compounds, at 0.45 ± 0.001% dw.

**Figure 2:**
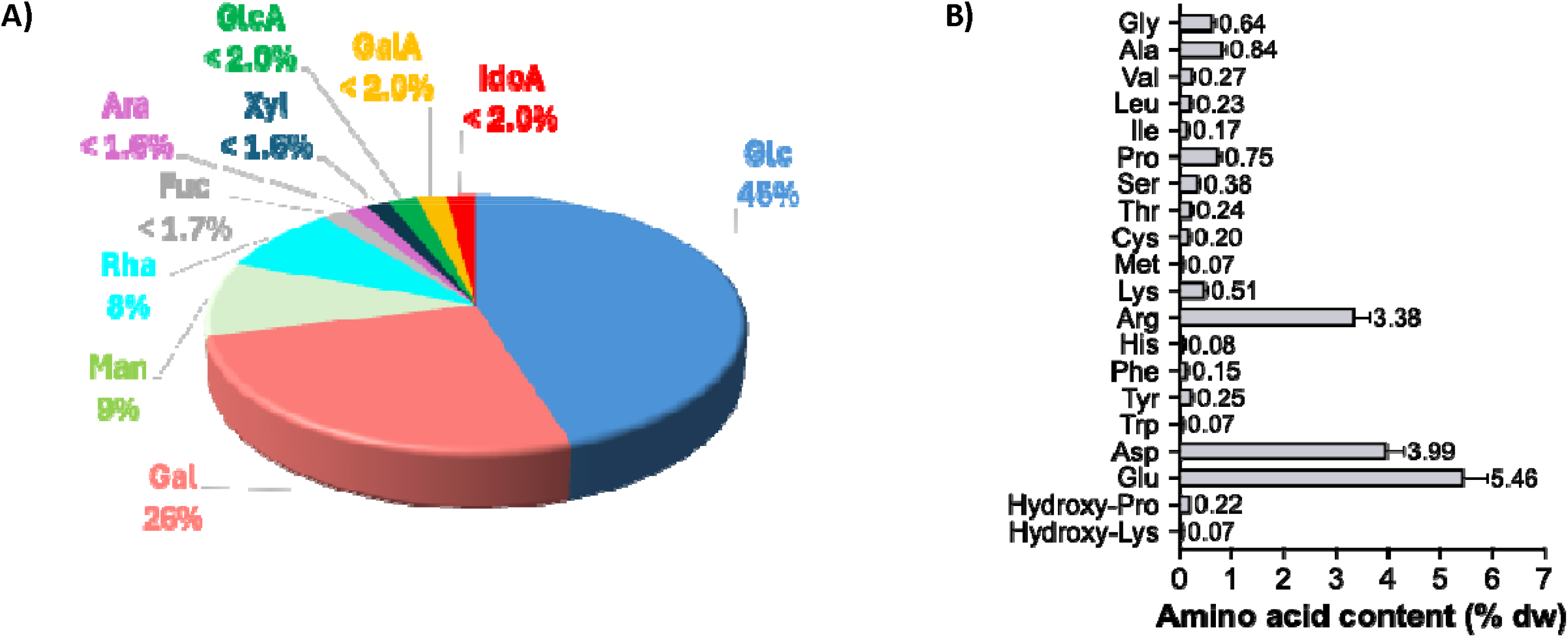
Osidic (A) and amino acid (B) compositions of permeate.

### Small organic compounds’ characterization of permeate

Given that permeate was composed as expected of a major part of low MW compounds (Figure 1), approximately half, in particular, at the small compound and/or oligomer scale (lower than 3 kDa), their nature was investigated using UHPLC-HRMS analysis. To increase both sensitivity and resolution of the analysis method, three different sample pre-treatment procedures were applied to obtain more simple and enriched fractions starting from permeate: 1) a size-based fractionation method that was also used to determine the weight distribution of permeate (Method 1; Figure S1); 2) three different forms of differential precipitation procedures associated to filtration methods (methods 2A, 2B and 2C; Figures S2, S3 and S4); and 3) two forms of acidic treatment using supported proton exchange. Twenty-one fractions were thus obtained and analyzed by untargeted MS^2^ analysis, leading to the identification of 53 organic compounds (Table 3), belonging to various biomolecule subclasses according to the ‘CANOPUS’ prediction module used through Sirius 4 data-processing, mostly amino acid-, fatty acid- and sugar-based structures. Amino acids, peptides, and analogues including 4 betaine derivatives (abromine, gynesine, stachydrine and beta-alanine betaine; entries 1, 5, 7, 23, respectively) and glutamine-containing small peptides (entries 2, 12, 34, respectively) were thus identified. Many amphiphilic fatty acids and conjugates, combining fatty acid chains to several unsaturations and/or polar functions such as hydroxylations and carbonyls were also found in permeate. Some of them were shown to belong to linoleic acid derivatives, carboxylic acids, or medium-chain hydroxy acids and derivatives subclasses, including several oxylipins (9-hydroxy-11-(3-hydroxy-5- (1-hydroxypropyl)tetrahydrofuran-2-yl)undec-10-enoic acid; phytoprostan A1; pinellic acid; fulgidic acid, 6,7,10-trihydroxy-8-octadecenoic acid; 6,7,12,13-tetrahydroxyoctadeca-8,10-dienoic acid; 6-hydroxyhexadeca-7,9,12,15-tetraenoic acid; 7-hydroxyoctadec-9-enedioic acid; and 9-oxooctadeca-10,12,15-trienoic acid: entries 11, 16, 20, 15, 29, 41, 42, 45, respectively), and low-chain fatty diacids (itaconate, suberate and azelaic acid: entries 6, 8, 3, respectively). Finally, some very small carbohydrate derivatives were detected in permeate: rhamnose-3-sulfate, one characteristic moiety composing the well-known primary structure of the so-called ulvans, parietal polysaccharides of *Ulva* genus seaweeds, and D-glucurono-6,3-lactone, trehalose and lilioside B (or C) (entries 21, 4, 22, 26). If some of these metabolites are well-known to be produced by *Ulva* genus seaweeds, such as abromine^47^, also called methyl- or glycine-betaine, which is a very common osmoprotectant in plants and microorganism, helping cells to resist osmotic stress (salt, drought) by stabilizing proteins and membranes^48^, some others had not been isolated from such biomass so far, or from marine macroalgae, to our knowledge: gynesine, stachydrine, beta-alanine betaine, oxylipins except phytoprostan A1^49^ and pinellic acid equivalents^50^, itaconate and suberate, D-glucurono-6,3-lactone, trehalose, lilioside B (or C). It is noteworthy that stachydrine, itaconate and D-glucurono-6,3-lactone were quantified in permeate at small but significant contents of 15.33 ± 1.19 and 7.40 ± 0.12 and 76.60 ± 14.47 mg/100 g dw.

**Table 3:**
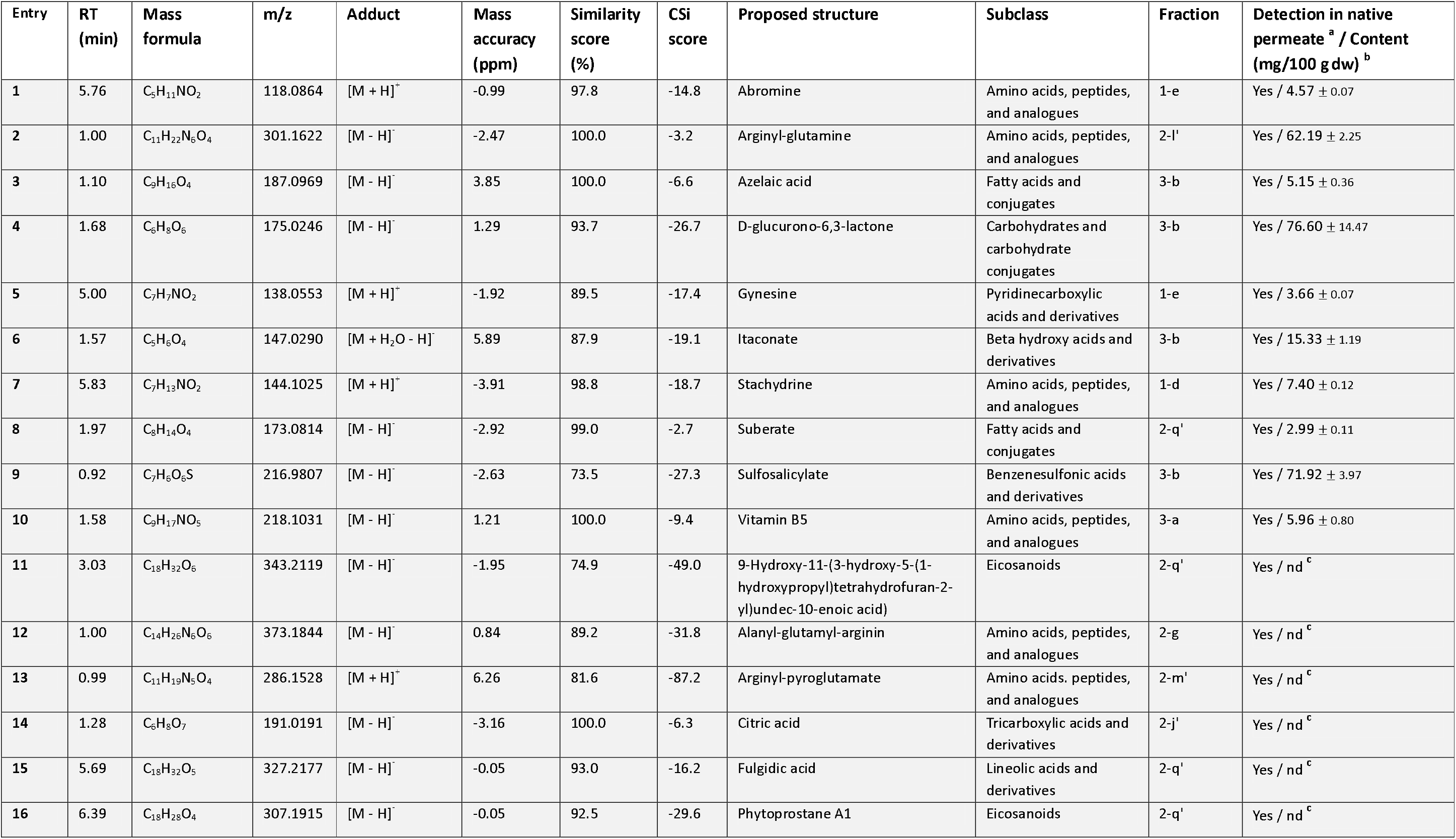
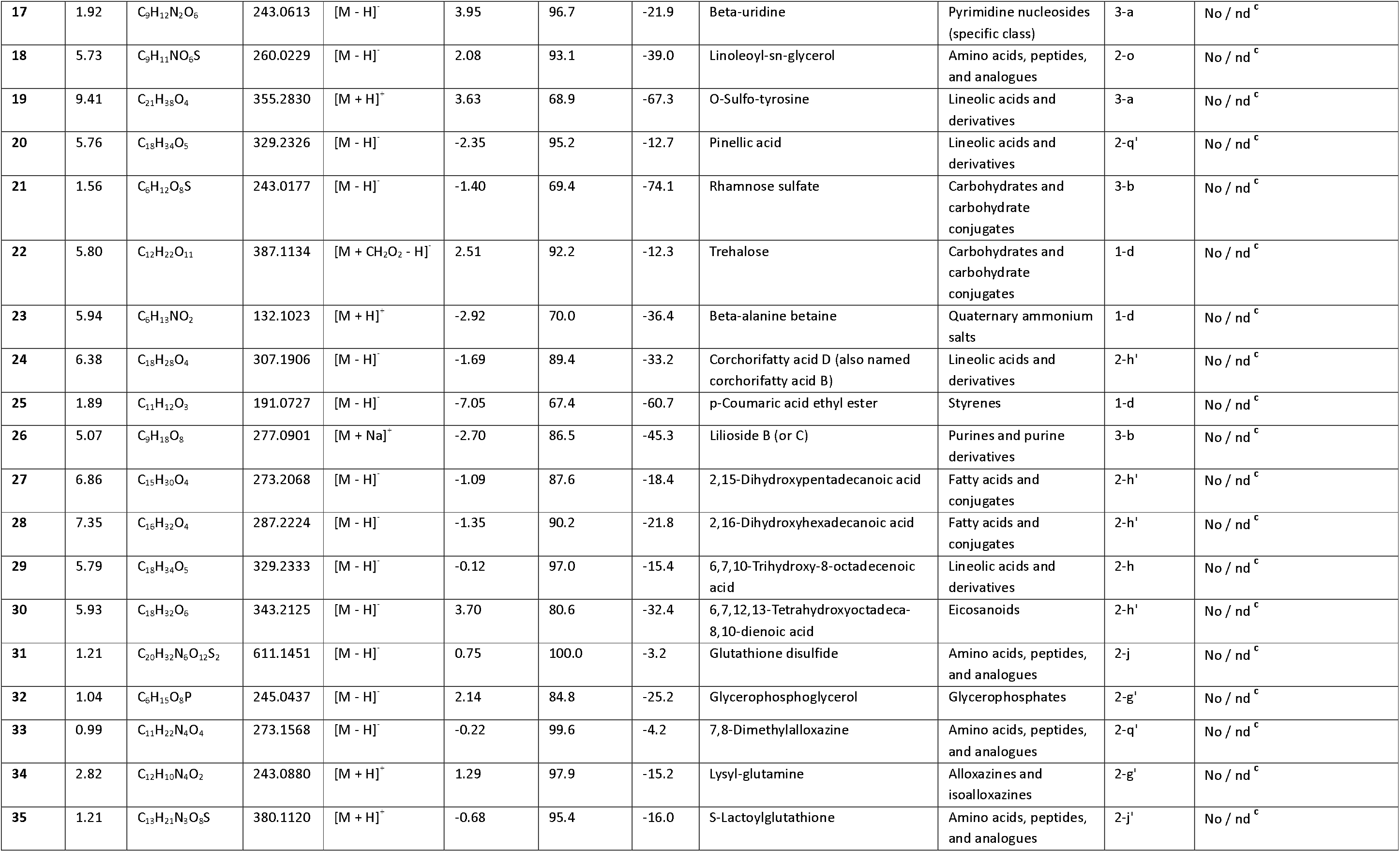
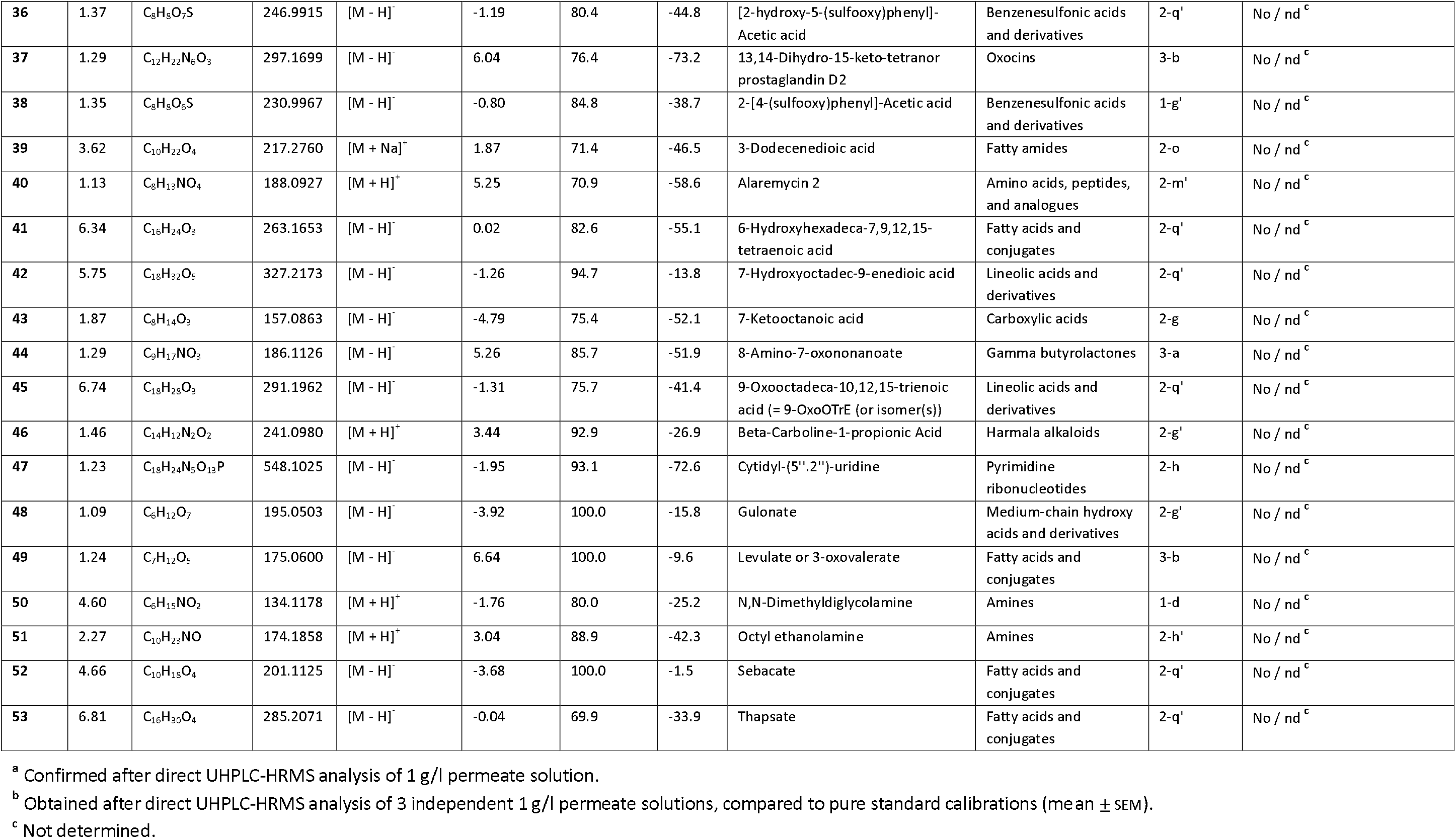
Compounds identified in permeate by UHPLC-HRMS analysis.

### Effect of permeate on RAW 264.7 macrophages’ viability

As a preliminary step, we checked the potential cytotoxicity of permeate towards macrophages (Figure 3). The results showed that permeate did not induce any detectable cytotoxic effects at tested concentrations, within the range 0.1-250 µg/ml. Based on these findings, lower concentrations, less or equal to 100 µg/ml, were selected for subsequent experiments, to test low potential active doses. In addition, LPS stimulation did not affect cell viability under the experimental conditions applied.

**Figure 3:**
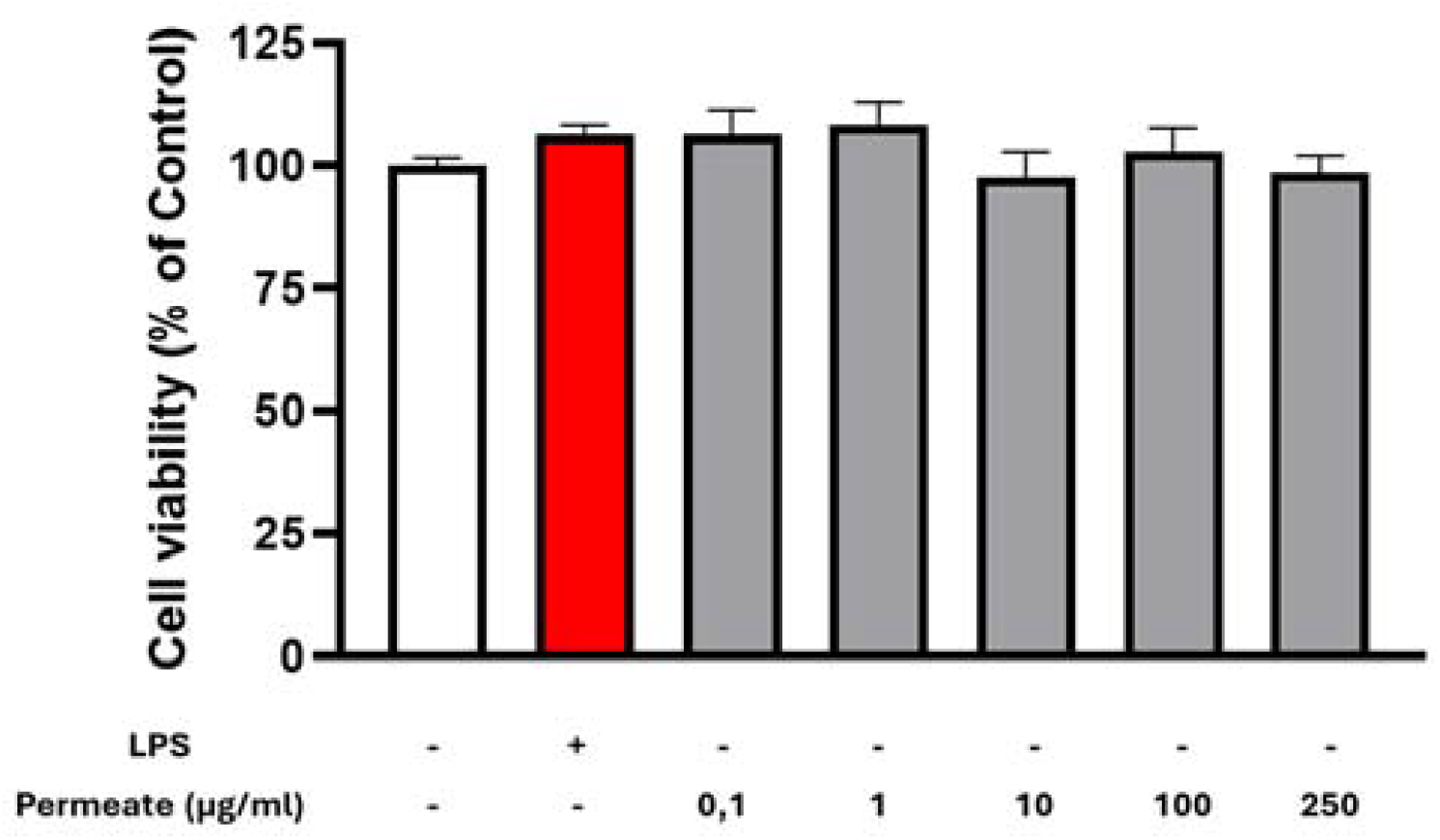
Effect of 24 h permeate exposition on RAW 264.7 macrophages’ viability *in vitro* (MTT assay). Results are expressed as the percentage of response compared to the control condition. n=4, ± SEM; LPS: 1µg/ml; p >0.05.

### Effect of permeate on pro-inflammatory cytokines’ gene expression in LPS-stimulated RAW 264.7 macrophages

The gene expression of the pro-inflammatory cytokines was evaluated in RAW 264.7 macrophages pre-treated with permeate for 24 h at 100 µg/ml prior to stimulation with LPS (1 µg/ml) for 2, 6 or 24 h. Permeate alone at 100 µg/ml did not significantly increase the expression of the pro-inflammatory cytokines IL-6, IL-1β and TNF-α compared with the control condition, except a modest increase in TNF-α at 2 h time point (p<0.01). Overall, it did not induce a pro-inflammatory response, at any time point. On the other hand, LPS stimulation induced a marked increase in TNF-α gene expression at all time points compared with unstimulated control cells (p<0.0001) (Figure 4A). Pre-treatment with permeate did not reduce TNF-α gene expression at early (2 h) or intermediate (6 h) time points but reduced it at 24 h (p<0.001), indicating a delayed modulatory effect. In contrast, pre-treatment with permeate significantly reduced IL-6 and IL-1β gene expression at all time points (p<0.001 and p<0.0001, respectively) (Figure 4B and C). These results indicate a sustained inhibitory effect of permeate on IL-6 and IL-1β transcriptional responses throughout the LPS stimulation period.

**Figure 4:**
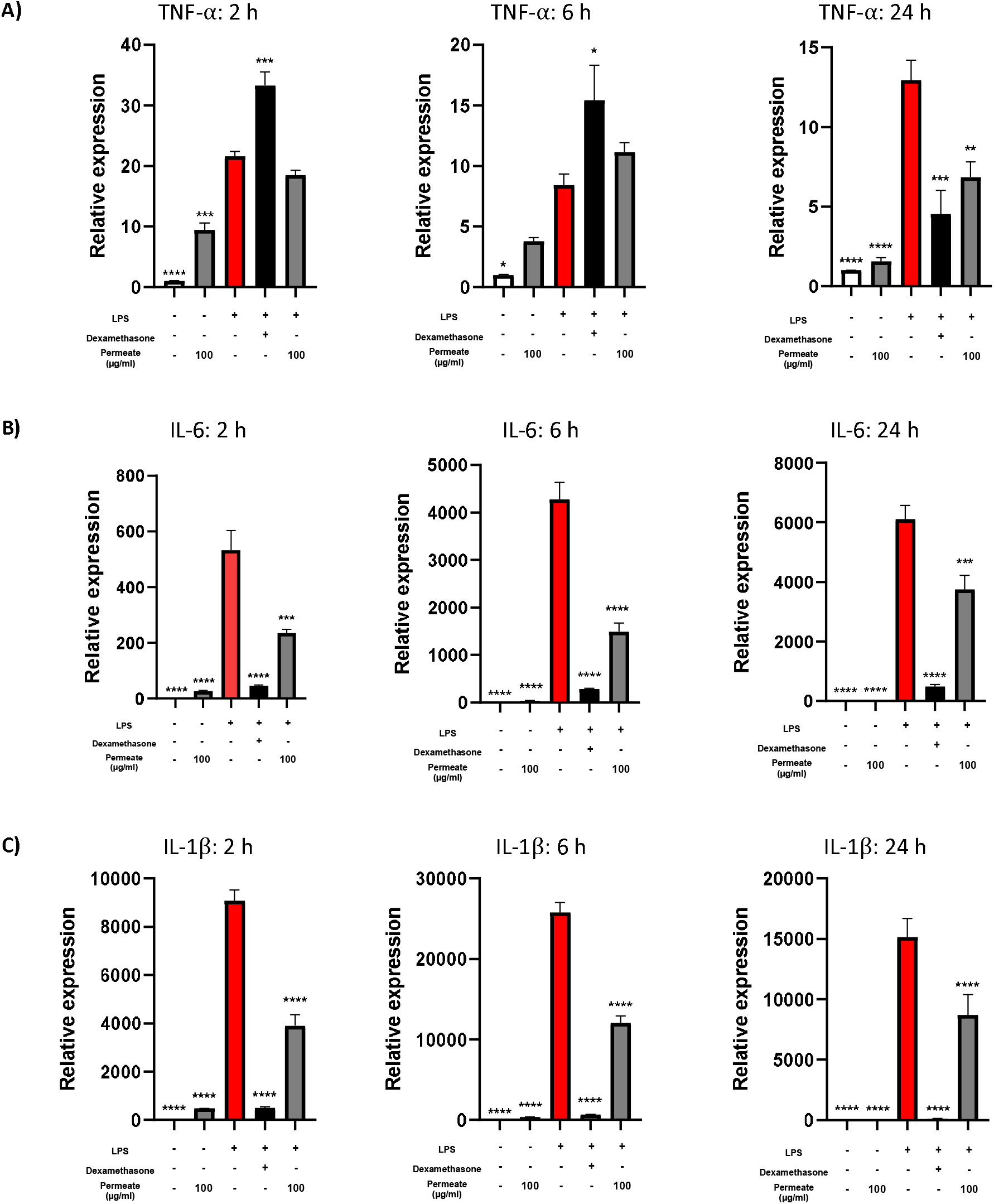
Effect of permeate pre-treatment on pro-inflammatory cytokines’ gene expression in LPS-stimulated RAW 264.7 macrophages *in vitro*. Macrophages were pre-incubated with 100 µg/ml permeate or 1 µM dexamethasone for 24 h and then incubated with both permeate or dexamethasone and 1 µg/ml LPS for the next 24 h. Cytokine expression of TNF-α (**A**), IL-6 (B) and IL-1β (**C**) was then measured by RT-qPCR. Results are expressed as the percentage of response compared to the positive control (LPS) condition. n=4, ± SEM; Dx: 1 µM dexamethasone; * p<0.05, ** p<0.01, *** p<0.001, **** p<0.0001.

### Effect of permeate on key pro-inflammatory mediators’ secretion in (LPS-stimulated) RAW 264.7 macrophages

RAW 264.7 macrophages incubated with permeate alone during 24 h did not trigger any pro-inflammatory response in comparison with control condition (Figure 5A, B and C), confirming the previous result obtained regarding pro-inflammatory cytokines’ gene expression. Across all tested concentrations, nitric oxide (NO) release remained indeed comparable to untreated control cells (Figure 5A). Similarly, the secretion of the two pro-inflammatory cytokines TNF-α and IL-6 (Figure 5B and C) was not significantly altered by permeate, with levels remaining low and within the range of unstimulated macrophages. On the contrary, LPS stimulation during 24 h logically induced strong secretions of the 3 pro-inflammatory cytokines (p<0.0001 for all).

**Figure 5:**
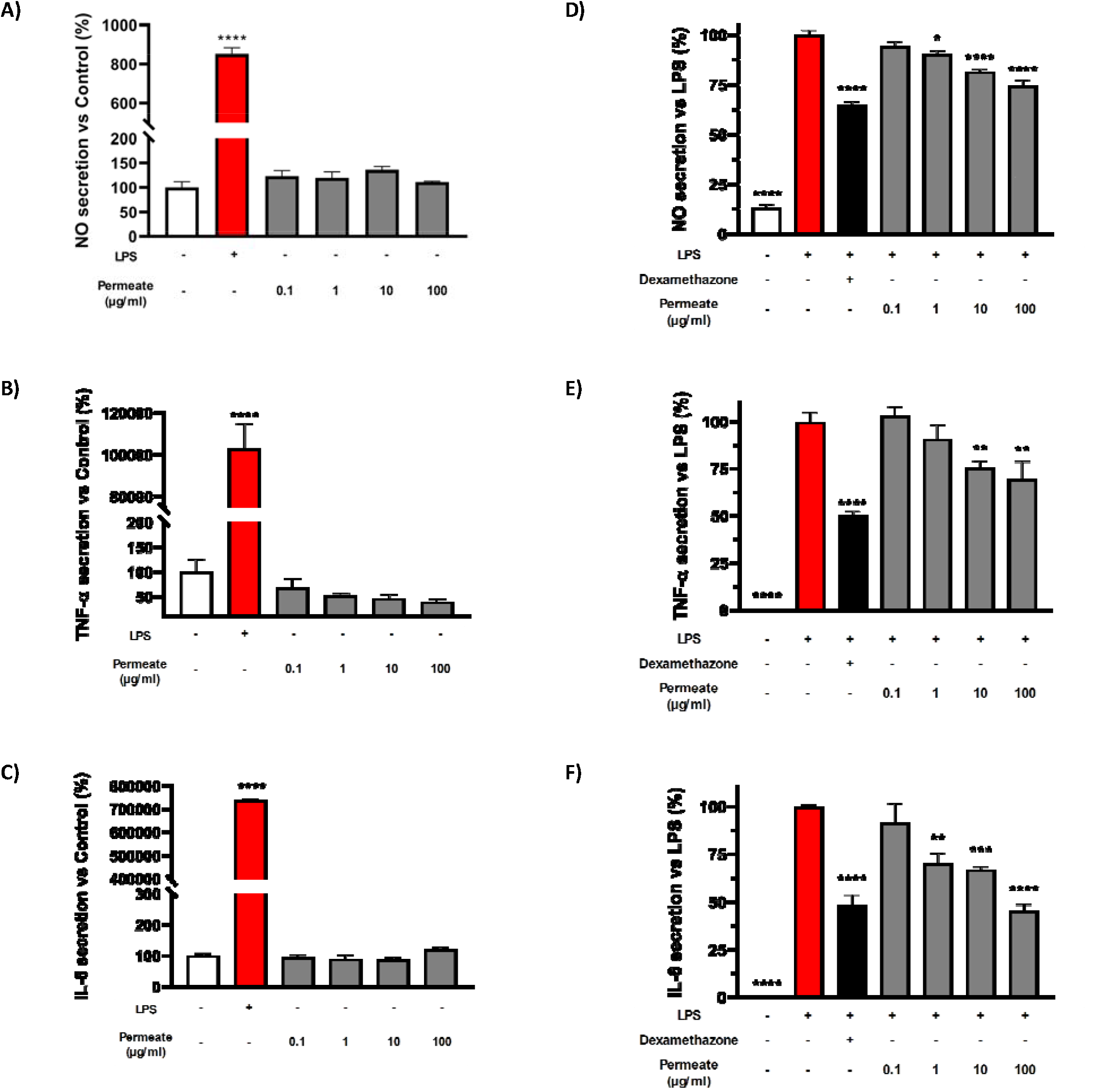
Effect of permeate pre-treatment on key pro-inflammatory mediators’ secretion in LPS-stimulated RAW 264.7 macrophages *in vitro*. Macrophages were incubated with permeate or 1 µg/ml LPS for 24 h (**A, B, C**), or pre-incubated with permeate or dexamethasone for 6 h and then incubated with both permeate or dexamethasone and LPS for the next 18 h (**D, E, F**). NO (**A, D**). TNF-α (**B, E**) and IL-6 (**C, F**) secretion was then measured. Results are expressed as the percentage of response compared to the control (**A, B, C**) or positive control (LPS) (**D, E, F**) condition. n=4, ± SEM; Dx: 1 µM dexamethasone; * p<0.05, ** p<0.01, *** p<0.001, **** p<0.0001.

Pre-treatment of macrophages during 6 h with permeate prior to 18 h of incubation with LPS significantly attenuated LPS-induced NO release in a concentration-dependent manner (Figure 5D). A statistically significant reduction in NO levels was observed starting from 1 µg/ml of permeate (p<0.0001), with a more pronounced secretion inhibition at higher concentrations. Similarly, IL-6 secretion was significantly decreased from 1 µg/ml (p<0.0001) onward (Figure 5F), indicating a strong sensitivity of this cytokine secretion to permeate pre-treatment. The reduction of TNF-α secretion required higher permeate concentrations, with a significant inhibitory effect observed from 10 µg/ml (p<0.0001) (Figure 5E). Importantly, the effect of permeate at 100 µg/ml was not significantly different from that of dexamethasone regarding IL-6 and TNF-α secretions (p-values of 0.9977 and 0.6958, respectively), indicating that this totum could be as potent as a pure anti-inflammatory compound.

### Modulation of inflammatory and cytoprotective proteins’ expression by permeate in LPS-stimulated macrophages

The effects of permeate on the expression of inflammation-related and cytoprotective proteins’ expression were evaluated in RAW 264.7 macrophages by western blot analyses. As expected, LPS stimulation induced a marked increase in COX-2 (p<0.0001) and NLRP3 (p<0.05) protein expression, as well as an increase in NF-κB phosphorylation (p<0.05), compared with untreated control cells (Figure 6A, B and C).

**Figure 6:**
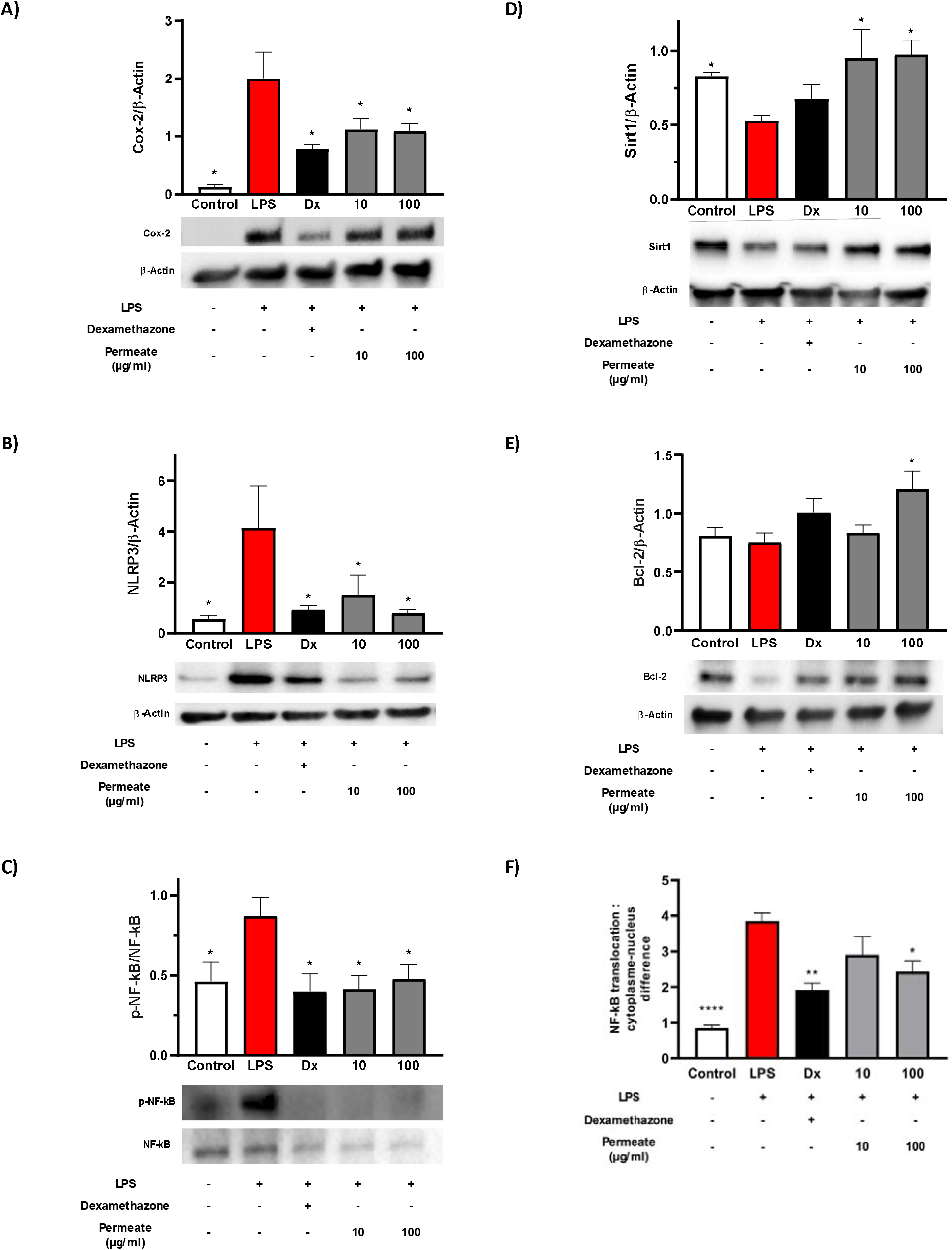
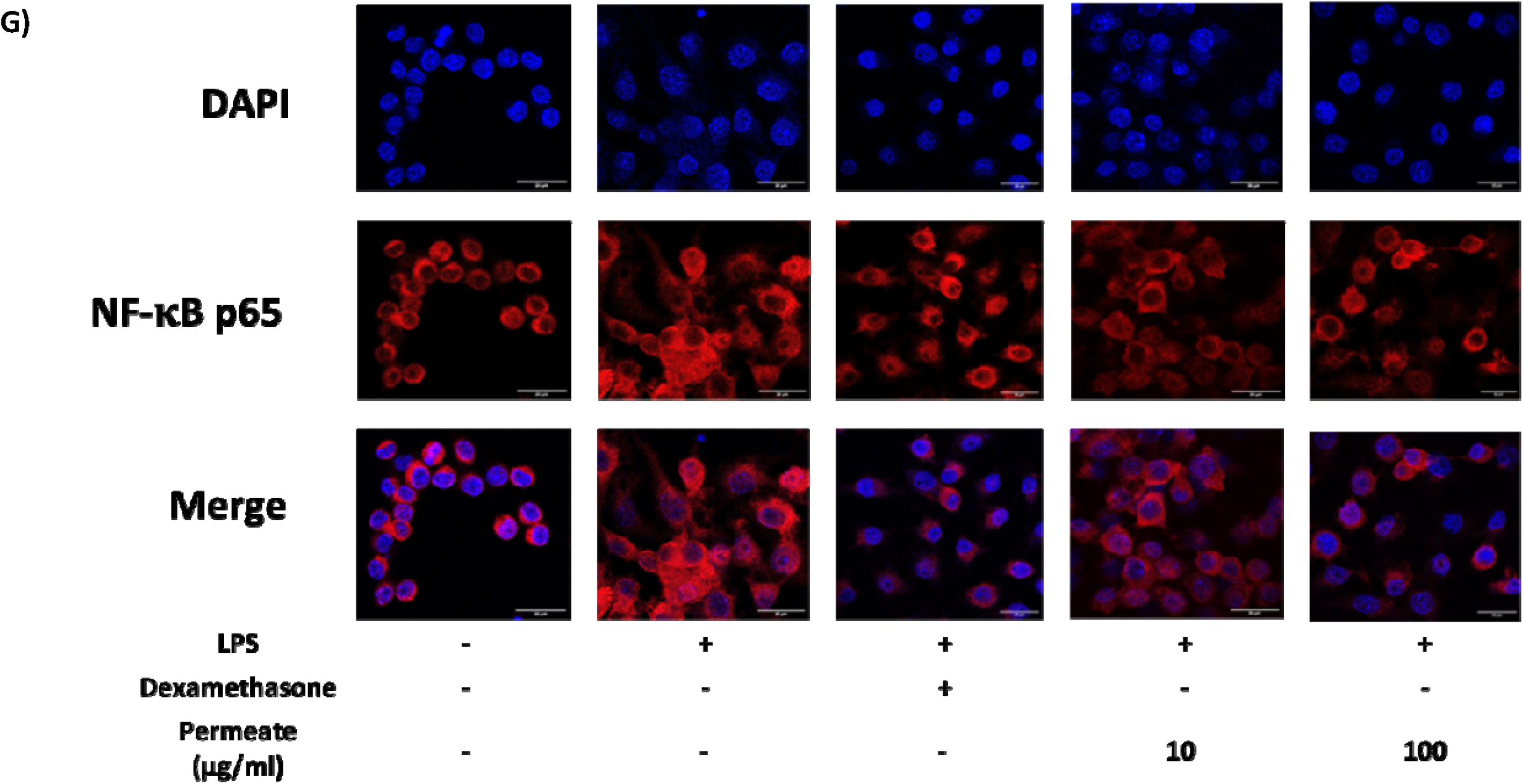
Effect of permeate pre-treatment on inflammatory and cytoprotective proteins’ expression in LPS-stimulated RAW 264.7 macrophages *in vitro*. Macrophages were pre-incubated with permeate or dexamethasone for 6 h and then incubated with both permeate or dexamethasone and 1 µg/ml LPS for the next 18 h. Protein expression was then measured by western-blot analysis and divided by the respective β-actin signal (**A-E**) or by total NF-kB (**F**), and cells were observed after fluorescent labelling by laser scanning confocal microscopy to determine the difference between cytoplasmic and nuclear NF-kB signals (**G**): the nuclear focal plane is reported on the pictures (grey scale bar = 10 μm). Results are expressed as the percentage of response compared to the positive control (LPS) condition. n=4, ± SEM; Dx: 1 µM dexamethasone; * p<0.05, ** p<0.01, *** p<0.001, **** p<0.0001.

Pre-treatment with dexamethasone markedly reduced LPS-induced NF-κB nuclear localization (p<0.001), confirming the validity of the experimental model. Pre-treatment with permeate reduced COX-2 and NLRP3 expression at both tested concentrations of 10 and 100 µg/ml, with lower protein expression levels observed in permeate-treated cells compared to LPS alone (p<0.05) (Figure 6A and B). In addition, a decrease in phosphorylated NF-κB levels was observed with permeate pre-treatment (p<0.05) (Figure 6C). Similarly to what was observed regarding IL-6 and TNF-α secretions, the effect of permeate was not significantly different from that of dexamethasone regarding COX-2 and NLRP3 expressions, as well as the phosphorylation of NF-κB, independently from its concentration: p-values of 0.5544, 0.9406 and 0.9999 at 10 µg/ml and 0.5925, 0.9998 and 0.9563 at 100 µg/ml, respectively. This confirmed the high potency of permeate as an anti-inflammatory ingredient.

Moreover, the protein expression of the cytoprotective proteins SIRT1 and Bcl-2 was significantly increased in permeate-treated cells compared with LPS-stimulated condition (p<0.05) (Figure 6D and E). This increase was observed at both concentrations tested regarding SIRT1, with a more pronounced effect at 100 µg/ml, but only at 100 µg/ml regarding Bcl-2. Using the same experimental conditions, the subcellular localization of NF-κB was examined by confocal microscopy in RAW 264.7 macrophages. As shown in Figure 6G, LPS stimulation induced a clear translocation of NF-κB from the cytoplasm to the nucleus, compared with untreated control cells (p<0.0001). Pre-treatment with permeate reduced NF-κB nuclear translocation in LPS-stimulated cells. Quantitative analysis of nuclear NF-κB signal intensity confirmed these observations as a significant decrease in NF-κB nuclear localization was shown under both dexamethasone (p<0.01) and 100 µg/ml permeate conditions (p<0.05), compared with LPS condition (Figure 6F). Together, these results indicate that permeate modulated NF-κB intracellular distribution under inflammatory conditions.

### Evaluation of autophagy-related signaling pathways in RAW 264.7 macrophages

To investigate whether permeate modulates autophagy, key regulators and markers of autophagy were analyzed in RAW 264.7 macrophages by Western blot. A 24 h-exposure of macrophages to 10 µg/ml permeate resulted in a significant dose-dependent increase in SIRT1 protein expression compared with untreated control cells (p<0.05) (Figure 7A). Analysis of AMP-activated protein kinase (AMPK) revealed that AMPK activation was affected by permeate (Figure 7B). The ratio of phosphorylated AMPK to total AMPK (p-AMPK/AMPK) was significantly increased in cells treated with 10 µg/ml permeate compared with control cells (p<0.05), indicating the activation of AMPK signaling. In contrast, no significant change in AMPK phosphorylation was detected at 100 µg/ml. Autophagy-related processing was further assessed by evaluating LC3 conversion (Figure 7C). A significant dose-dependent increase in the LC3-II/LC3-I ratio was observed in cells treated with 10 µg/ml (p<0.05) and 100 µg/ml (p<0.01) permeate, compared with control conditions, suggesting enhanced autophagosome formation. In addition to molecular markers, cell membrane and cytoskeleton labelling of macrophages with phalloidin and DAPI, respectively, was examined by confocal microscopy (Figure 7D). Permeate-treated macrophages displayed a marked morphological change compared with control cells, tending towards a much more dendritic form associated to a significant increase in their relative volume at 100 g/ml permeate (p<0.05) (Figure 7E), indicating an actin cytoskeleton reorganization typical of autophagy-associated morphological changes.

**Figure 7:**
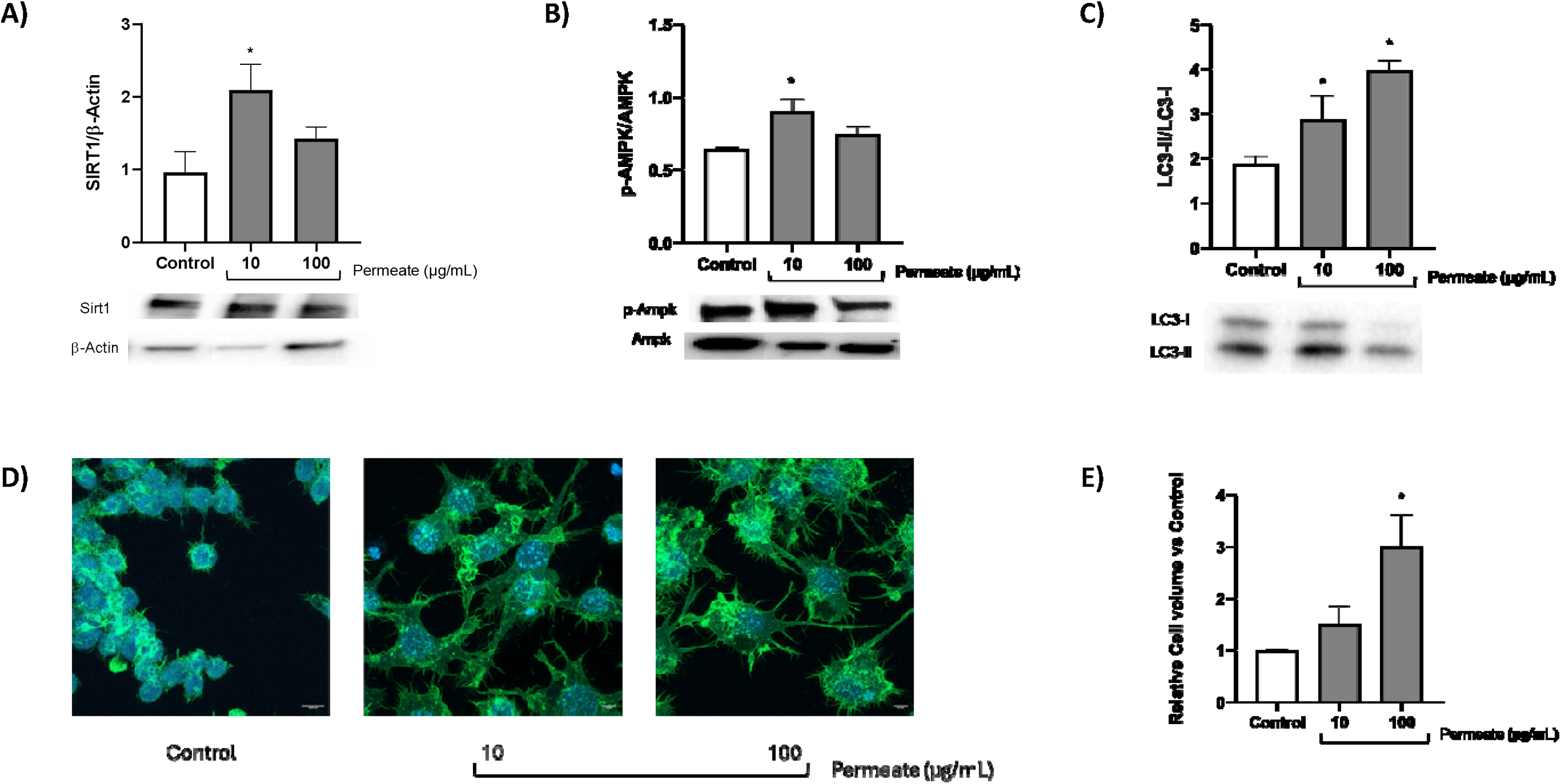
Effect of permeate treatment on key autophagy markers’ protein expression in RAW 264.7 macrophages *in vitro*. Macrophages were incubated with 10 or 100 µg/ml permeate for 24 h. Protein expression of SIRT1 (**A**; rapported to β-Actin signal), p-AMPK (**B**; rapported to AMPK total expression) and LC3-II (**C**; rapported to LC3-I), was then measured by western-blot analysis and cells were observed after specific nuclear and cytoskeleton fluorescent labelling by laser scanning confocal microscopy (**D**; the nuclear focal plane is reported on the pictures: grey scale bar = 10 μm), allowing to measure their relative cell volume (**E**; graphical representation of cytoplasmic area (fluorescent Alexa Fluor 594 dye-conjugated phalloidin; green) divided by the number of nuclei detected in the corresponding field (DAPI dye; blue)). Results are expressed as the percentage of response compared to the positive control (LPS) condition. n=3, ± SEM; * p<0.05.

## Discussion

This study provides a comprehensive evaluation of a low MW compounds rich-permeate derived from *Ulva lacinulata*, combining detailed chemical characterization with functional investigations in inflammatory macrophage models. Due to the use of a patented ultrafiltration process with a 15 kDa cut-off, permeate was rich in small compounds, about 50% of less than 3 kDa. This was of particular interest from the nutritional point of view as it was shown to contain minerals of nutritional interest, especially the macro elements magnesium (3265.5 ± 42.0 g/100 g dw) and sulfur (10204.4 ± 114.0 g/100 g dw), and the trace elements iodine (3.20 ± 0.64 g /100 g dw) and manganese (17.96 ± 0.05 g/100 g dw), as the consumption of 1 gram of permeate per day would bring about 11%, 17%, 21% and 9% of their recommended nutrient intake, respectively^45^; oligo- and poly-saccharides involving ulvan and starch structures; or proteins and therefore amino acids, mostly arginine, aspartic acid glutamic acid and, to a lesser extent, the essential amino acids Val, Lys, Phe and Thr. It was also proved that permeate contained very little quantities of heavy metals (17 metals analyzed), largely below the limits accepted in food^46^, and did not contain any phytosanitary product (336 pesticides analyzed and undetected, data not shown), which allows to consider its further potential application as an active safe ingredient for functional food and/or dietary supplements.

As a preliminary step regarding the *in vitro* activity of permeate, we showed that it did not induce any detectable cytotoxic effect towards macrophages at all tested concentrations (within the range 0.1-250 µg/ml). This was in accordance with the absence of cytokine secretion induced by permeate, suggesting that the extract did not trigger cellular stress responses that could secondarily promote inflammation as described in the literature by Fulda et al.^51^. Furthermore, these findings strongly support the effectiveness of the fractionation process and, more specifically, of the 15 kDa ultrafiltration cut-off applied to eliminate high–MW biopolymers, especially polysaccharides such as ulvans, which have been described in the literature as pro-inflammatory and/or immunomodulatory when present in large molecular forms^5–7^.

We next investigated NO, TNF-α, IL-1β, IL-6, Cox-2 and NLRP3 expression and/or secretion by LPS-induced RAW 264.7 macrophages, as a relevant and widely accepted approach for assessing *in vitro* inflammatory activation^52,53^. Indeed, upon LPS stimulation, macrophages rapidly upregulate inducible inflammatory pathways leading to the increased NO production via inducible nitric oxide synthase (iNOS) expression, as well as the secretion of pro-inflammatory cytokines such as TNF-α, IL-1β and IL-6^54^. Permeate significantly reduced the expression and/or secretion of these key pro-inflammatory mediators in LPS-stimulated macrophages in a dose-dependent manner, strongly supporting the biological relevance of the observed effects. Furthermore, the magnitude and direction of these inhibitory effects were supported by the result obtained with dexamethasone. From a mechanistic perspective, the concomitant reduction of NO, TNF-α, IL-1β and IL-6 suggests that permeate interfered with upstream inflammatory signaling events common to these mediators, rather than targeting a single isolated pathway. These results were further supported by the phosphorylation and subcellular localization of NF-κB that were significantly decreased by 100 µg/ml permeate pre-treatment. Indeed, NF-κB is a central transcriptional regulator of pro-inflammatory genes, including those encoding iNOS, TNF-α, IL-1β and IL-6. The attenuation of its phosphorylation thus provides a mechanistic explanation for the coordinated decrease in these inflammatory mediators^55,56^. Besides, NF-κB requires phosphorylation to dissociate from its cytoplasmic inhibitors and translocate into the nucleus, where it promotes the transcription initiation of pro-inflammatory genes^57^. This finding provides complementary spatial evidence that permeate interfered with NF-κB activation at an early stage of the inflammatory cascade.

Moreover, permeate pre-treated cells significantly reduced the expression of downstream inflammatory effectors, including Cox-2 and the NLRP3 inflammasome component, when submitted to LPS stimulation. Cox-2 is a well-established NF-κB–regulated enzyme involved in the synthesis of pro-inflammatory prostaglandins^58^, while NLRP3 plays a central role in inflammasome assembly and the amplification of inflammatory signaling^59^. The concomitant downregulation of Cox-2 and NLRP3 expression further supports the hypothesis that permeate exerted a broad inhibitory effect on macrophage inflammatory activation.

These anti-inflammatory effects observed in LPS-stimulated RAW 264.7 macrophages in our study, characterized by a significant reduction in nitric oxide secretion and both gene expression and secretion of TNF-α and IL-6, can be mechanistically linked to the diverse set of bioactive metabolites identified in the algal permeate. Importantly, treatment with permeate alone did not induce a pro-inflammatory response, indicating that its activity was not immunomodulatory per se but rather modulatory in the context of inflammatory challenge. This profile is consistent with the presence of compounds that selectively interfere with inflammatory signaling pathways activated upon LPS stimulation. Indeed, a substantial proportion of the metabolites detected in permeate have been reported to inhibit the NF-κB signaling cascade, which plays a central role in macrophage activation and pro-inflammatory gene expression. Compounds such as stachydrine^60^, azelaic acid^61^, phytoprostane A1^62^, pinellic acid^63^ and citric acid^64^ have been shown to suppress NF-κB activation or nuclear translocation, resulting in reduced transcription of iNOS, TNF-α and IL-6. The coordinated inhibition of this pathway by multiple constituents likely underlies the marked attenuation of inflammatory mediator production observed in our results and may explain the dose-dependent nature of permeate effects. Emerging evidence also highlights the importance of macrophage metabolic state in shaping inflammatory responses^65^. Metabolites such as itaconate^66^, which was quantified at a dose of 15.33 ± 1.19 mg/100 g dw in permeate, and beta-uridine^67^, also identified in permeate, have been implicated in immuno-metabolic reprogramming toward an anti-inflammatory phenotype. In particular, itaconate has been shown to suppress cytokine production by inhibiting inflammasome activation and promoting Nrf2-dependent transcriptional programs in macrophages^68^. The presence of such metabolite supports the hypothesis that permeate did not merely block inflammatory signaling but could also actively promote a metabolic environment unfavorable to sustained cytokine secretion in macrophages^69^.

Lipid-derived metabolites constitute another key component of permeate anti-inflammatory profile. Oxylipins, hydroxylated fatty acids and monoacylglycerols, including phytoprostanes such as phytoprostane A1^70^, phytofurans such as (E)-9-hydroxy-11-(3-hydroxy-5-(1-hydroxypropyl)tetrahydrofuran-2-yl)undec-10-enoic acid)^70^, pinellic acid^71^ and linoleoyl glycerol, are known modulators of lipid-mediated inflammatory signaling. These compounds can interfere with membrane-associated receptor signaling and the synthesis of pro-inflammatory lipid mediators, thereby contributing to the observed decrease in cytokine and NO release. Their combined presence further emphasizes the multi-layered regulation of inflammation exerted by permeate. Finally, peptides, glycosides and sulfated compounds identified in permeate, such as arginyl-glutamine^72^, alanyl-glutamyl-arginine^73,74^ and lilioside B/C^75^, have been reported to modulate macrophage signaling and cytokine secretion. While their individual effects may be moderate, their cumulative action may likely contribute to the fine-tuning of macrophage responsiveness to inflammatory stimuli.

Together, these findings indicate that the anti-inflammatory activity of *Ulva lacinulata* permeate most likely arises from the synergistic action of multiple metabolites targeting complementary inflammatory pathways, rather than from a single dominant compound. To further elucidate the mechanism of action of permeate, we investigated its effects under basal conditions prior to inflammatory stimulation, focusing on molecular markers involved in immuno-metabolic regulation and autophagy. Studying these basal responses is critical to distinguish adaptive cellular signaling from secondary effects associated with LPS-induced inflammation. This approach allows a clearer interpretation of whether permeate primes macrophages toward a protective phenotype rather than merely counteracting LPS-induced damage.

Autophagy has been increasingly recognized as a key cellular mechanism contributing to the resolution of inflammation, particularly in macrophages, where it participates in the degradation of pro-inflammatory signaling components and the restoration of metabolic homeostasis^76,77^. Beyond its canonical role in cellular quality control, autophagy tightly interacts with innate immune pathways, including NF-κB and inflammasome signaling^78,79^. Importantly, pharmacological or nutritional modulation of autophagy has emerged as a promising strategy to limit excessive inflammatory responses while preserving essential immune functions^80^. Permeate exposure induced a coordinated modulation of SIRT1 expression, AMPK phosphorylation, and autophagy-related marker LC3. Such coordinated regulation suggests the engagement of a conserved metabolic sensing network rather than isolated signaling events. This pattern is consistent with a controlled cellular adaptation aimed at optimizing energy balance and limiting pro-inflammatory activation^81,82^. An increase in AMPK phosphorylation was indeed observed in macrophages treated with 10 µg/ml permeate, consistent with the activation of an early energy-sensing pathway known to promote autophagy initiation and to negatively regulate inflammatory signaling. AMPK activation has also been reported to suppress NF-κB-dependent transcription^83^. Consistent with upstream metabolic signaling, permeate exposure also induced a concentration-dependent modulation of the autophagic pathway. After 24 h exposure to permeate, SIRT1 expression and AMPK phosphorylation were increased at 10 µg/ml permeate but not at 100 µg/ml. This apparent discrepancy might be explained by the occurrence of a faster resolution of the initial activation phase of the autophagy pathway at the highest concentration of permeate, leading to the decline in upstream autophagy regulators such as SIRT1 and p-AMPK. It might also be due to the complexity of permeate which, as a totum, could associate both synergical and antagonistic phenomena occurring at different concentrations. On the contrary, LC3-II/LC3-I ratio was increased in a dose-dependent manner. LC3-I lipidation leading to LC3-II formation represents a downstream event in the autophagic cascade^84^, which suggests that autophagosome formation has already occurred and the autophagic process is still resolving after 24 h exposure to permeate. Autophagy is indeed a dynamic and sequential process, in which upstream metabolic sensors and downstream structural markers are temporally uncoupled. AMPK and SIRT1 regulate different phases of autophagy initiation and progression, whereas LC3-I lipidation and autophagosome maturation occur later during a complete autophagic cycle^85^. Such coordinated dynamics are characteristic of controlled, non-pathological autophagy^86^.

Several metabolites identified in permeate have been previously reported to activate these autophagy-related pathways. For instance, itaconate^87^ and trehalose^88^ are known inducers of autophagy: itaconate promotes mitophagy through Nrf2 activation and metabolic reprogramming, while trehalose activates autophagy via an mTOR-independent mechanism, consistent with the increased LC3-II levels observed in our study. Similarly, azelaic acid^89^ and stachydrine^60^ have been shown to stimulate AMPK signaling, which in turn can activate SIRT1 and enhance autophagic flux, providing a likely mechanistic explanation for the coordinated increase in SIRT1 expression and AMPK phosphorylation observed in permeate-treated macrophages. The activation of SIRT1 by these metabolites may also contribute to NF-κB inhibition induced by permeate, possibly linking autophagy induction to the reduction of LPS-induced pro-inflammatory cytokines (NO, TNF-α, IL-6). Indeed, SIRT1 is well-known to deacetylate the p65 sub-unit of NF-κB, limiting its transcriptional activity, while AMPK activation promotes autophagosome formation and metabolic adaptation^90^. Altogether, these results suggest the involvement of an autophagy-mediated effect rather than isolated inhibition of inflammatory pathways induced by permeate. By engaging the SIRT1–AMPK–autophagy axis, permeate seems to establish a cellular state that favors resolution over activation, enabling macrophages to mount a restrained inflammatory response when challenged by LPS. Such an indirect but robust mechanism is particularly relevant in the context of chronic low-grade inflammation, where long-term modulation of immune responsiveness is preferable to acute immunosuppression^91^. Furthermore, autophagy has been extensively described as a critical regulator of inflammasome activity, particularly through the selective removal of damaged mitochondria and the suppression of mitochondrial danger signals that drive NLRP3 activation, such as excessive mitochondrial reactive oxygen species (ROS) production, cytosolic release of oxidized mitochondrial DNA, cardiolipin exposure, and ATP-mediated purinergic signaling^92,93^. By promoting autophagic flux and therefore autophagy-mediated cellular housekeeping, permeate may limit the accumulation of mitochondrial-derived activators and inflammatory intermediates required for NLRP3 inflammasome assembly, leading to the reduced production of NLRP3 observed in permeate-treated macrophages (Figure 6B). However, further investigations are still needed to ascertain this assumption and precisely understand to what extent the stimulation of autophagy by permeate accounts for its anti-inflammatory activity, especially under inflammation conditions induced by LPS.

In conclusion, the present study demonstrated that the *Ulva lacinulata* extract permeate exerts a robust anti-inflammatory activity in cultured RAW 264.7 macrophages, as evidenced by the coordinated reduction of NO production and pro-inflammatory cytokine expression and secretion, together with the inhibition of NF-κB signaling. Importantly, these effects occurred without triggering basal inflammatory activation or affecting macrophage viability, supporting the safety and potential biological compatibility of permeate. Beyond cytokine suppression, permeate modulated key immuno-metabolic regulators such as SIRT1 and AMPK and engaged the autophagic machinery in a controlled manner, as reflected by LC3 activation. This coordinated regulation suggests that permeate did not merely counteract inflammatory insults but also promoted a cellular state conducive to metabolic resilience and immune homeostasis. Such mechanisms are increasingly recognized as central to the regulation of low-grade inflammation, a systemic condition strongly implicated in functional decline, across various pathophysiological contexts such as aging and overtraining.

Taken together, these findings provide a strong rationale for considering this *Ulva lacinulata* extract permeate as a promising candidate for the development of an active ingredient for functional food and/or dietary supplements targeting chronic low-grade inflammation. By acting on fundamental inflammatory and immuno-metabolic pathways in macrophages, permeate may contribute to improving systemic inflammatory balance through nutritional intervention. We are currently conducting large scale in vivo studies to validate these effects in various physiological contexts, to assess bioavailability and safety of permeate, and to determine whether long-term dietary supplementation with permeate may mitigate peripheral and/or brain inflammation and, directly or indirectly, limiting pathways leading to neurodegeneration. At last, some of the identified metabolites in permeate such as beta-alanine betaine, itaconate and D-glucurono-6,3-lactone are well-known redox-regulating compounds that might further support autophagy by reducing oxidative stress, which is both a known suppressor of autophagic flux and a strong multi-levels activator of inflammation. The potential effect of the *Ulva lacinulata* extract permeate on oxidative stress of LPS-induced RAW 264.7 macrophages is therefore also under ongoing investigation by our team.

## Acknowledgement

We thank Seprosys for providing the *Ulva lacinulata* permeate. We thank the “High resolution analysis of biomolecules” platform from LIENSs laboratory. We are grateful to C. Churlaud and M. Brault-Favrou from the “Elemental Analysis” platform of LIENSs laboratory for their assistance during trace element analyses. We thank the NutriBrain Research and Technology Transfer platform of NutriNeuro laboratory. We thank the “Centre d’Etude et de Valorisation des Algues” (CEVA). We also thank the Alg4Health consortium.

