## Supplemental Figure S1 for "Low-molecular-weight *Ulva lacinulata* extract exhibiting anti-inflammatory and pro-autophagic activities in RAW 264.7 macrophages: a promising candidate for the development of active ingredients targeting low-grade inflammation"

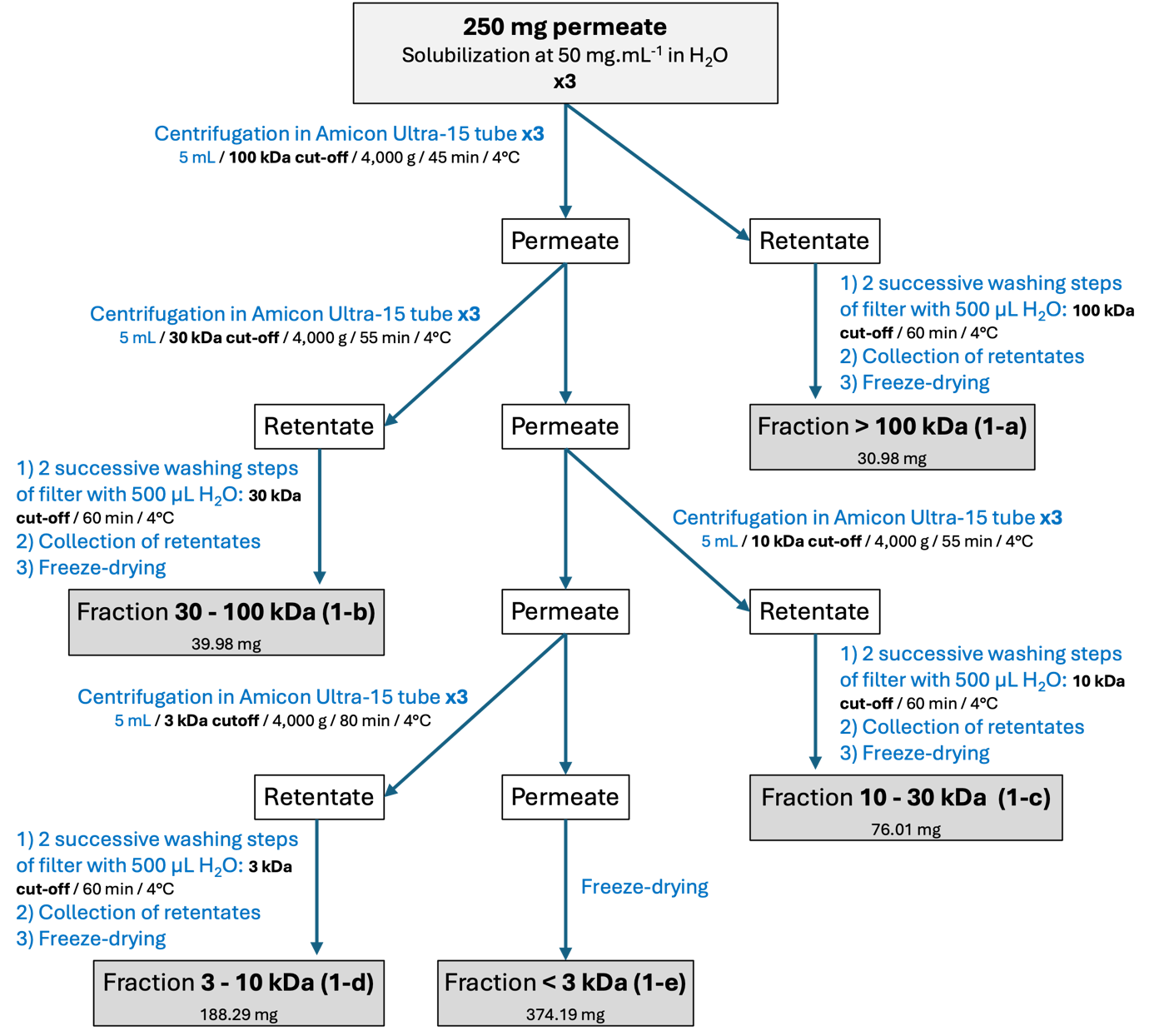

**Figure S1: Method 1 of size-based fractionation used to both determine the weight distribution of permeate and to produce permeate fractions 1a-1e.**
