## Supplemental Figure S3 for "Low-molecular-weight *Ulva lacinulata* extract exhibiting anti-inflammatory and pro-autophagic activities in RAW 264.7 macrophages: a promising candidate for the development of active ingredients targeting low-grade inflammation"

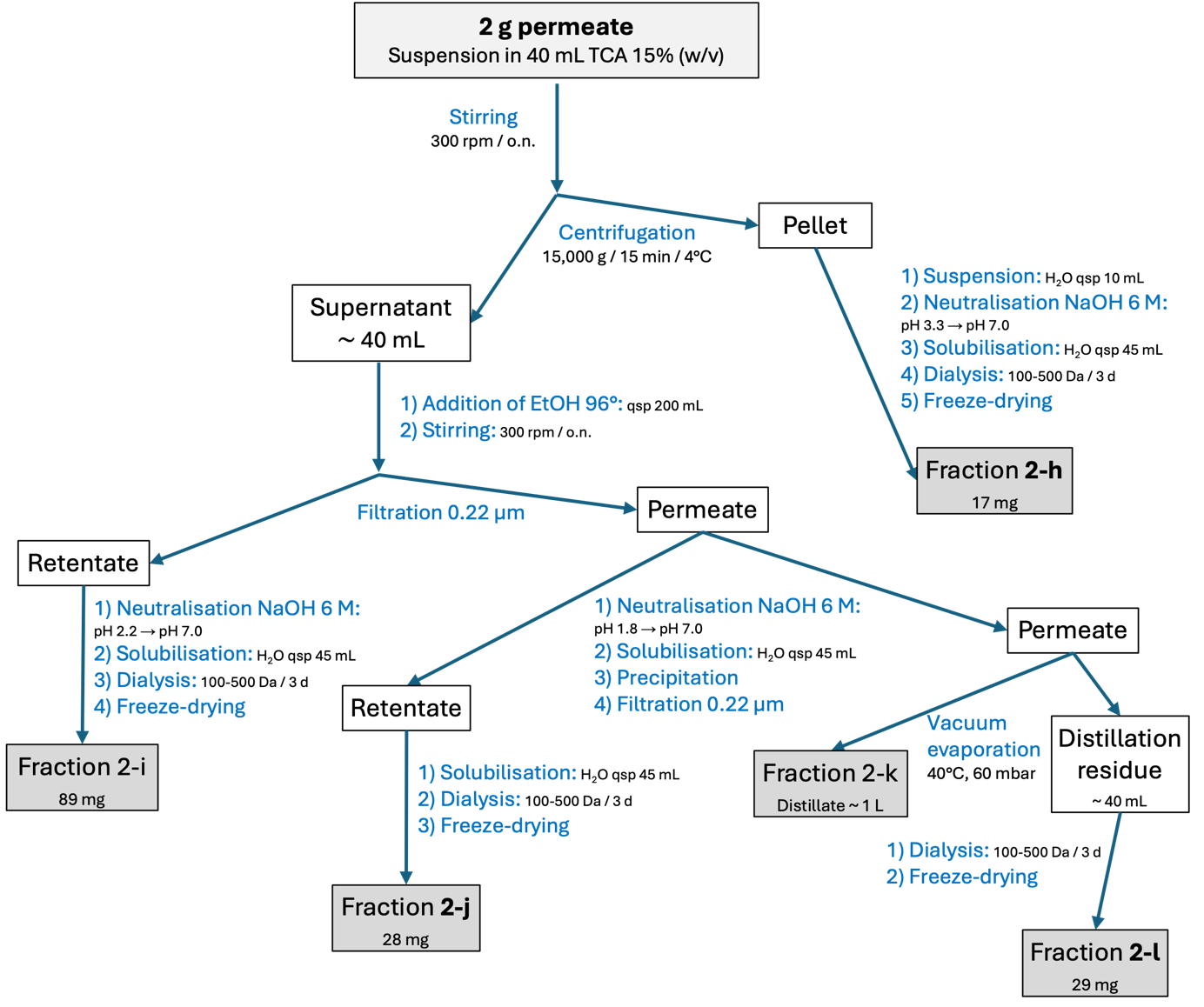

**Figure S3: Method 2B of differential precipitation associated to filtration steps used to obtain permeate fractions 2h-2l.**
