## Supplemental Figure S4 for "Low-molecular-weight *Ulva lacinulata* extract exhibiting anti-inflammatory and pro-autophagic activities in RAW 264.7 macrophages: a promising candidate for the development of active ingredients targeting low-grade inflammation"

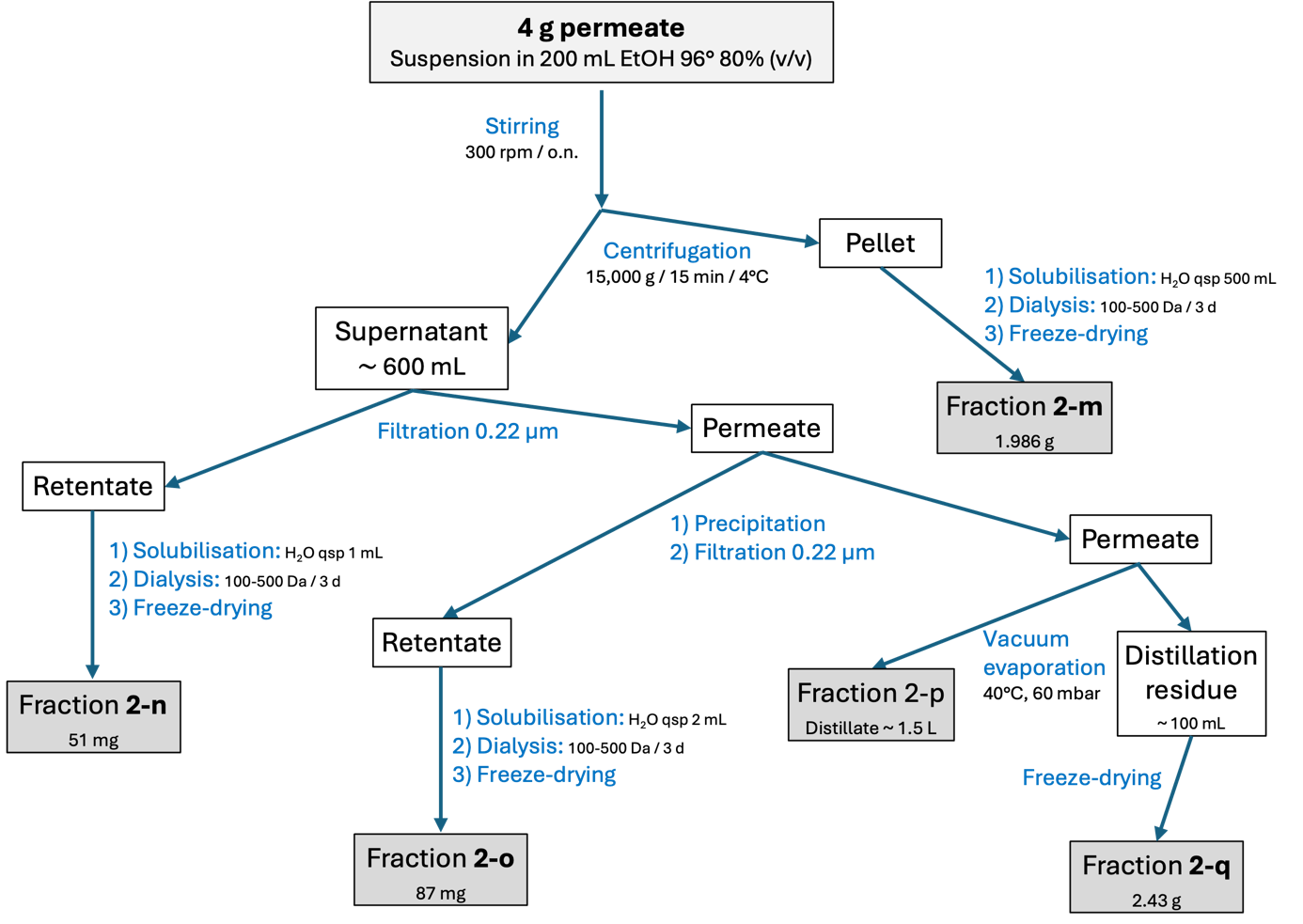

**Figure S4: Method 2C of differential precipitation associated to filtration steps used to obtain permeate fractions 2m-2q.**
